# Extensive hybridization between pig and human *Ascaris* identifies a highly interbred species complex infecting humans

**DOI:** 10.1101/2020.04.17.047407

**Authors:** Alice V. Easton, Shenghan Gao, Scott P Lawton, Sasisekhar Bennuru, Asis Khan, Eric Dahlstrom, Rita G Oliveira, Stella Kepha, Steve F Porcella, Joanne P Webster, Roy M Anderson, Michael E. Grigg, Richard E Davis, Jianbin Wang, Thomas B Nutman

**Affiliations:** Helminth Immunology Section, Laboratory of Parasitic Diseases, National Institute of Allergy and Infectious Disease, National Institutes of Health, Bethesda, Maryland 20892, USA; Department of Infectious Disease Epidemiology, Imperial College London, London W2 1PG, UK; Department of Biochemistry and Molecular Genetics, RNA Bioscience Initiative, University of Colorado School of Medicine, Aurora, Colorado 80045, USA; Beijing Institute of Genomics, Chinese Academy of Sciences, Beijing 100101, China; Molecular Parasitology Laboratory, School of Life Sciences, Pharmacy & Chemistry, Kingston University London, Kingston Upon Thames, Surrey KT1 2EE, UK; Molecular Parasitology Section, Laboratory of Parasitic Diseases, National Institute of Allergy and Infectious Disease, National Institutes of Health, Bethesda, Maryland 20892, USA; Genomics Unit, Research Technologies Branch, National Institute of Allergy and Infectious Diseases, National Institutes of Health, Hamilton, MT 59840, USA; London School of Tropical Medicine and Hygiene, Keppel St, Bloomsbury, London, WCIE 7HT, UK; Royal Veterinary College, University of London, Department of Pathobiology and Population Sciences, Hertfordshire AL9 7TA, UK; Department of Biochemistry and Cellular and Molecular Biology, University of Tennessee, Knoxville, Tennessee 37996, USA

## Abstract

Human ascariasis is a major neglected tropical disease caused by the nematode *Ascaris lumbricoides*. We report a 296 megabase (Mb) reference quality genome comprised of 17902 protein-coding genes derived from a single, representative *Ascaris* worm collected from 60 human hosts in Kenyan villages where pig husbandry is rare. Notably, the majority of human isolates (63/68) possessed mitochondrial genomes that clustered closer to the pig parasite *Ascaris suum* than to *A. lumbricoides*. Comparative phylogenomic analyses identified over 11 million nuclear-encoded SNPs but just two distinct genetic types that had recombined across the genomes analysed. The nuclear genomes had extensive heterozygosity and all samples existed as genetic mosaics with either *A. suum*-like or *A. lumbricoides*-like inheritance patterns supporting a highly interbred *Ascaris* species genetic complex. As no barriers appear to exist for anthroponotic transmission of these “hybrid” worms, a one-health approach to control the spread of human ascariasis will be necessary.

## Introduction

Approximately 447 million people were estimated to be infected with the intestinal nematode *Ascaris lumbricoides* in 2017, resulting in an estimated 3206 deaths and a loss of over 860,000 Disability-Adjusted Life Years (DALYs, Global Burden of Disease Study 2017 http://ghdx.healthdata.org/gbd-2017). Many infections go undiagnosed, but like other soil-transmitted helminths (STH), *Ascaris* spp. infections contribute significantly to global DALYs, perpetuating the cycle of poverty in areas of endemic infection ^1–4^. Despite the large global burden of STH, little is known about *A. lumbricoides* transmission patterns or the true prevalence of infection with the pig parasite *A. suum* infection in people in endemic regions.

Deworming has become more widespread in areas of endemic STH infection ^5^. Regional health authorities and global health organizations are now looking for strategies to build on these programs by achieving local elimination of STH as a public health problem ^6^. A greater understanding of transmission dynamics (including the frequency of zoonotic transmission) using molecular epidemiological methods in settings where *A. lumbricoides* prevalence is low but persistent could help move current efforts towards successfully eliminating transmission through more targeted treatment.

Population genetic studies of *A. lumbricoides* have drawn varying conclusions about whether zoonotic transmission is frequent ^7–10^. Some studies have shown that cross-species transmission occurs between pigs and humans living in close proximity ^9,11–16^. This is especially common in non-endemic regions, probably because zoonotic transmission is less likely to be identified in areas where human-to-human transmission is common. To date, published results fail to conclude whether the human parasite *A. lumbricoides* and the pig parasite *Ascaris suum* are capable of interbreeding, and it is generally accepted that they exist as separate species. Furthermore, it is unclear whether pigs are an important reservoir of infection in humans worldwide or if *A. suum* is readily transmitted anthroponotically ^7,17–20^. Studies have generally concluded that the genetic differences between *Ascaris* worms collected from human populations in different parts of the world ^11,21^ are the result of geographic reproductive isolation. Previous studies using *Ascaris* mitochondrial genomes or genes suggest there are *A. lumbricoides*-type (human-associated) and *A. suum*-type (pig-associated) clades ^8,22,23^. Other work suggests multiple clades of worms, only one of which is unique to pigs ^24^.

In the current study, we constructed a reference-quality *Ascaris* genome (ALV5) based on sequences from a single female worm collected from a single person in Kenya. This person was presumed to be infected with *A. lumbricoides* as there is a lack of local pig husbandry. Draft *A. suum* genomes have previously been constructed from worms obtained from pigs in Australia ^25^ and in the United States ^26,27^. The *Ascaris* genome ALV5 was found to be highly similar (99% identity) to the *A. suum* genome from worms collected from pigs in the United States ^28^. Our mitochondrial and whole-genome analyses from an additional 68 individual worms indicate that *A. suum* and *A. lumbricoides* form a genetic complex that is capable of interbreeding. Our data support a model for a recent worldwide, multi-species *Ascaris* population expansion caused by the movement of humans and/or livestock globally. *Ascaris* isolates from both pigs and humans may be important in human disease, necessitating a one-health approach to control the spread of human ascariasis.

## Results

### Human *Ascaris* reference genome to promote comparative genomic analyses

To generate a human *Ascaris* spp. germline genome assembly (prior to programmed DNA elimination ^28^), ovarian DNA was sequenced from a single female worm collected from a Kenyan who was presumed to be infected with *A. lumbricoides* using Illumina paired-end and mate-pair libraries of various insert sizes with a total sequence coverage of ∼27-fold (Table S1). Using these data, three different assembly strategies were used (see below and supplemental information).

The *de novo* assembly and semi-*de novo* strategies produced poor *A. lumbricoides* germline draft genomes (Table 1 and supplementary text). In the semi-*de novo* assembly, the majority of the >4000 short contigs (making up 15.4 Mb of sequence) that could not be incorporated into the semi-*de novo* assembly are sequences that aligned to the genome at multiple positions. Comparison of the *A. suum* gene annotations to this assembly revealed a low *A. lumbricoides* gene number and high numbers of partial and split genes (Table 1, see footnote 3). These characteristics are typical of highly fragmented genomes or genomes with high levels of mis-assemblies ^28^.

**Table 1.** *Ascaris* germline genome assemblies.

| Features | *A. lumbricoides*<br>*de novo* | *A. lumbricoides*<br>semi-*de novo*<sup>1</sup> | *A. lumbricoides*<br>reference-based | *A. suum*<sup>2</sup><br>(Wang et al 2017) | *A. suum*<sup>2</sup><br>(Jex et al 2011)<sup>4</sup> |
| --- | --- | --- | --- | --- | --- |
| Assembled bases (Mb) | 269.2 | 307.9 | 296.0 | 298.0 | 272.8 |
| N50 (Mb) | 0.29 | 4.77 | 4.63 | 4.65 | 0.41 |
| N50 number | 269 | 21 | 21 | 21 | 179 |
| N90 (Mb) | 0.04 | 0.95 | 0.91 | 0.92 | 0.08 |
| N90 number | 1,112 | 74 | 75 | 75 | 748 |
| Total scaffold number | 8,111 | 412 | 415 | 415 | 29,831 |
| Largest scaffold length (Mb) | 1.9 | 13.9 | 13.2 | 13.4 | 3.8 |
| Protein-coding genes | 17,011<sup>3</sup> | 17,105<sup>3</sup> | 17902 | 18,025 | 18,542<sup>3</sup> |

<sup>1</sup> Exhibits ~23 Mb of sequence gaps and 15.4 Mb of unplaced sequence in 4,072 short contigs.
<sup>2</sup> The three *A. lumbricoides* assemblies constructed here are compared to the *A. suum* assemblies from Australia<sup>25</sup> and the United States<sup>28</sup>.
<sup>3</sup> 21-23% are only partial genes based on the annotation from *A. suum*<sup>28</sup>.
<sup>4</sup> The sample for sequencing is derived from a mixture of the germline and somatic genomes (after DNA elimination).

Mapping of the human *Ascaris* reads to the *A. suum* reference genome ^28^ revealed an exceptionally high sequence similarity (>99% identity) between the two species with few human *Ascaris* reads that could not be mapped to *A. suum*. Based on this high sequence similarity, a third reference-based-only assembly strategy was used to generate the human *Ascaris* germline genome assembly using the *A. suum* germline genome as a reference (see methods). This approach led to a reference-quality human *Ascaris* genome assembly with many fewer gaps (only 0.98 Mb of sequence) and no unplaced contigs. The *Ascaris* genome assembled into 415 scaffolds with a combined size of 296 Mb. An additional 15.4 Mb of sequence was present in 4072 unscaffolded short contigs. The assembly N50 value was 4.63 Mb, with the largest scaffold measuring 13.2 Mb. The largest 50 scaffolds combined to represent 78% of the genome. The assembly was further polished using additional Illumina reads from the same worm to more accurately reflect single base differences, indels, and any potential local mis-assembled regions.

To evaluate the quality of the assembled genome, we mapped the *Ascaris* Illumina reads back to the reference-based *Ascaris* genome assembly and found that > 99% of the Illumina reads could be mapped, indicating that the reference-based assembly excluded very few *Ascaris* reads. We then mapped and transferred the extensive set of *A. suum* transcripts ^25,28^ to the human *Ascaris* germline assembly to annotate the genome, identifying and classifying 17902 protein-coding genes (Table 1, Table S2). As this reference-based assembly exhibits the best assembly attributes, including high continuity with a large N50, low gaps and unplaced sequences, and high-quality protein-coding genes (see Table 1), we suggest that this version should be used as a reference germline genome for a human *Ascaris* spp. isolate (available in NCBI GenBank with accession number PRJNA515325). The other two assemblies are available and are discussed in more detail in the supplemental text.

Like *A. suum* embryos, *A. lumbricoides* embryos undergo programmed DNA elimination during the differentiation of the somatic cells from the germline in early development ^29,30^. In *A. suum*, ∼30 Mb of 120 bp tandem repeats and ∼1000 germline-expressed genes are lost from the germline to form the somatic genome ^27,28^. We also sequenced the somatic genome from the intestine of the same female *A. lumbricoides* worm. Comparison of the germline and somatic genomes revealed that DNA elimination in the human *Ascaris* isolate (including the breaks, sequences, and genes eliminated) was identical to that described for the pig *A. suum* isolate ^28^.

### Gene content and *Ascaris* proteome

Earlier annotations of protein coding genes for *A. suum* draft genomes were produced by Jex ^25^ and Wang ^27^ and improved with a recent updated genome ^28^—though the focus of the recent study was not on protein annotations. Here, we updated, identified, and fully annotated the 17902 protein-coding genes in the reference-based genome assembly (Table S2 and Figure S1), an annotation that can be a resource for others studying *Ascaris*. Using a custom pipeline (see methods and ^31^), we classified 48% of the predicted proteome into functional groups (Figure 1A). Although the remaining 52% (9300) of the genes were classified as unknown/uncharacterized, 2515 (27%) of these appear to encode proteins that have signatures indicative of either being secreted or being membrane-bound (some with GPI anchors). To provide a more comprehensive annotation of the transcriptomes of *A. suum* and *A. lumbricoides*, we re-mapped the RNA-seq data from *A. suum* to the current gene models of *A. lumbricoides* (ALV5) (Table S2). We performed multivariate analyses of this revised RNA-seq data compilation to generate a comprehensive RNA-seq data set for differential gene expression in diverse stages/tissues (Table S2).

**Figure 1.**
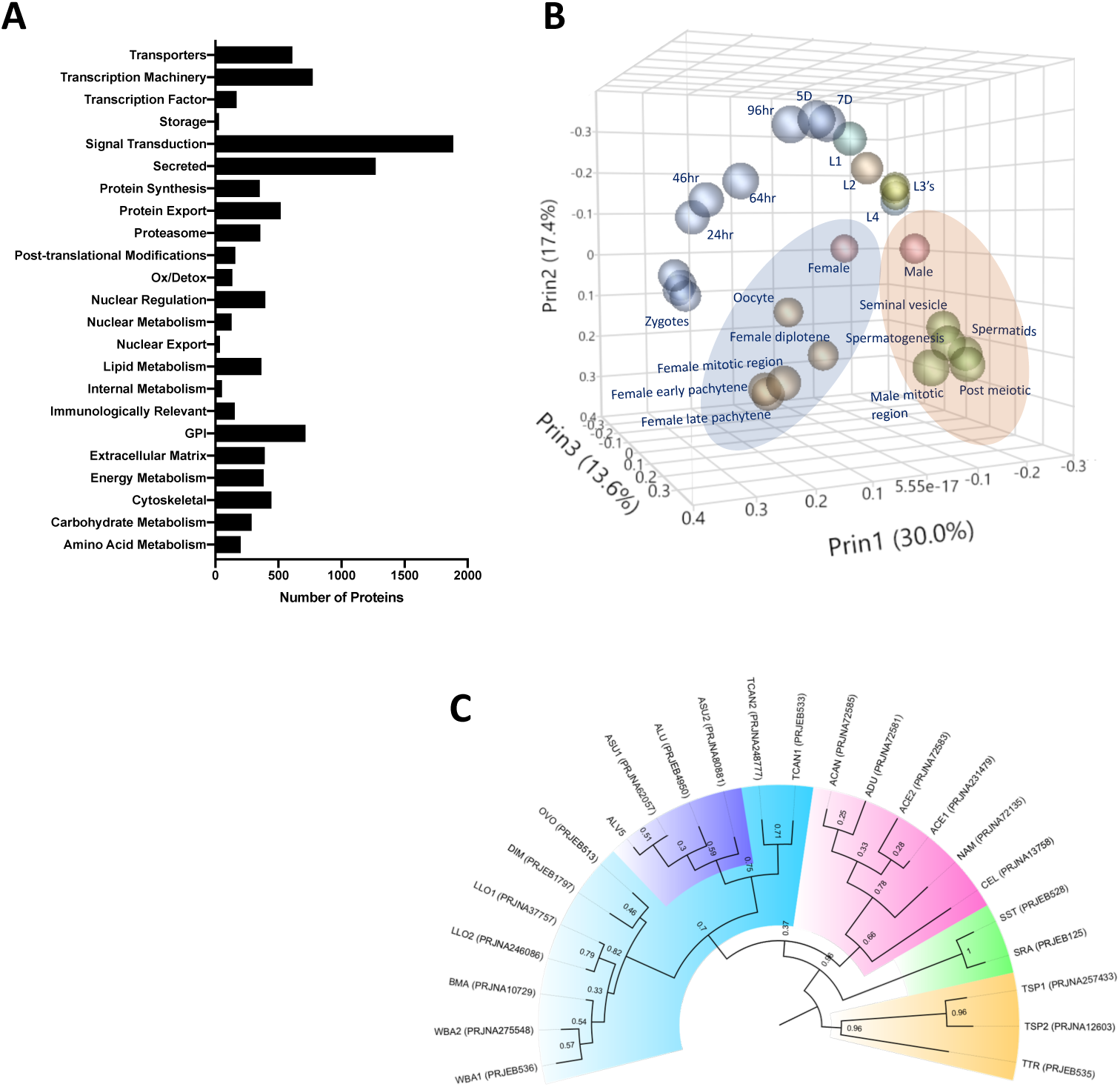
*Ascaris* proteome. A) Functional classification of the predicted proteome of *A. lumbricoides* (an improved proteome of *Ascaris* spp.), excluding proteins with unknown or uncharacterized function. B) PCA plot based on multivariate analyses of RNA-seq data from various stages/tissues. Samples from tissues related to sperm (blue ellipse) and oocyte production (orange ellipse, see also Figure S9) cluster together. C) Estimated tree based on orthology analyses between the predicted proteomes of publicly available nematodes. The *Ascaris* clade has been shaded in purple within Clade III (teal). Samples are labelled by BioProject Accession number, as well as by the first letter of the genus and the first two letters of the species name (ASU = *Ascaris suum*, ALU = *Ascaris lumbricoides*, WBA = *Wuchereria bancrofti*, BMA = *Brugia malayi*, LLO = *Loa loa*, DIM = *Dirofilaria immitis*, OVO = *Onchocerca volvulus*, TCAN = *Toxocara canis*, ACAN = *Ancylostoma caninum*, ADU = *Ancylostoma duodenale*, ACE = *Ancylostoma ceylanicum*, NAM = *Necator americanus*, CEL = *Caenorhabditis elegans*, SST = *Strongyloides stercoralis*, SRA = *Strongyloides ratti*, TSP = *Trichinella spiralis*, TTR = *Trichuris trichiura*). Multiple genomes for the same organism are suffixed with numerals.

Phylogenetic trees derived from orthologue analyses of the predicted proteomes of ALV5 with the predicted proteomes of other nematodes across all clades indicated the similarity among the published genomes of *A. suum* [PRJNA62057 and PRJNA80881 in ^25,27,28^] and *A. lumbricoides* ^32^ with ALV5 within the *Ascaris* branch (Figure 1C). The variation observed within the *Ascaris* spp. (with relatively weak bootstrap values of 0.3-0.59) is likely due to the differences in protein coding gene annotations and split genes seen in previous assemblies.

### Mitochondrial genome assembly

We next took advantage of the abundant reads from the mitochondrial genome in our sequencing data (on average 7690X coverage, see Table S1) to perform *de novo* assembly of 68 complete human isolate *Ascaris* spp. mitochondrial genomes from individual worms (Table S3). These mitochondrial genomes were then annotated using sequence similarity to well-characterized and annotated mitochondrial genes.

### Population structure inferred from mitochondrial CO1 gene

The mitochondrial CO1 gene has been frequently used to infer evolutionary distances between species as well as between populations ^22,33–37^ due to its rapid mutation rate, lack of recombination and relatively constant rate of change over time ^38–40^. Previous CO1 phylogeny studies resolve *Ascaris* spp. worms into three distinct clades: clade A is predominantly comprised of worms isolated from pigs, clade B is predominantly comprised of worms isolated from humans, and clade C is from worms only isolated from pigs in Europe and Asia ^22^. Interestingly, haplotype network analyses revealed that the majority of worms isolated from humans in the Kenyan villages possessed CO1 haplotypes that were consistent with infection of parasites from clade A (63/68), whereas only 6 isolates had CO1 haplotypes consistent with infection by worms from clade B (Figure S2 and Figure 2a)).

**Figure 2.**
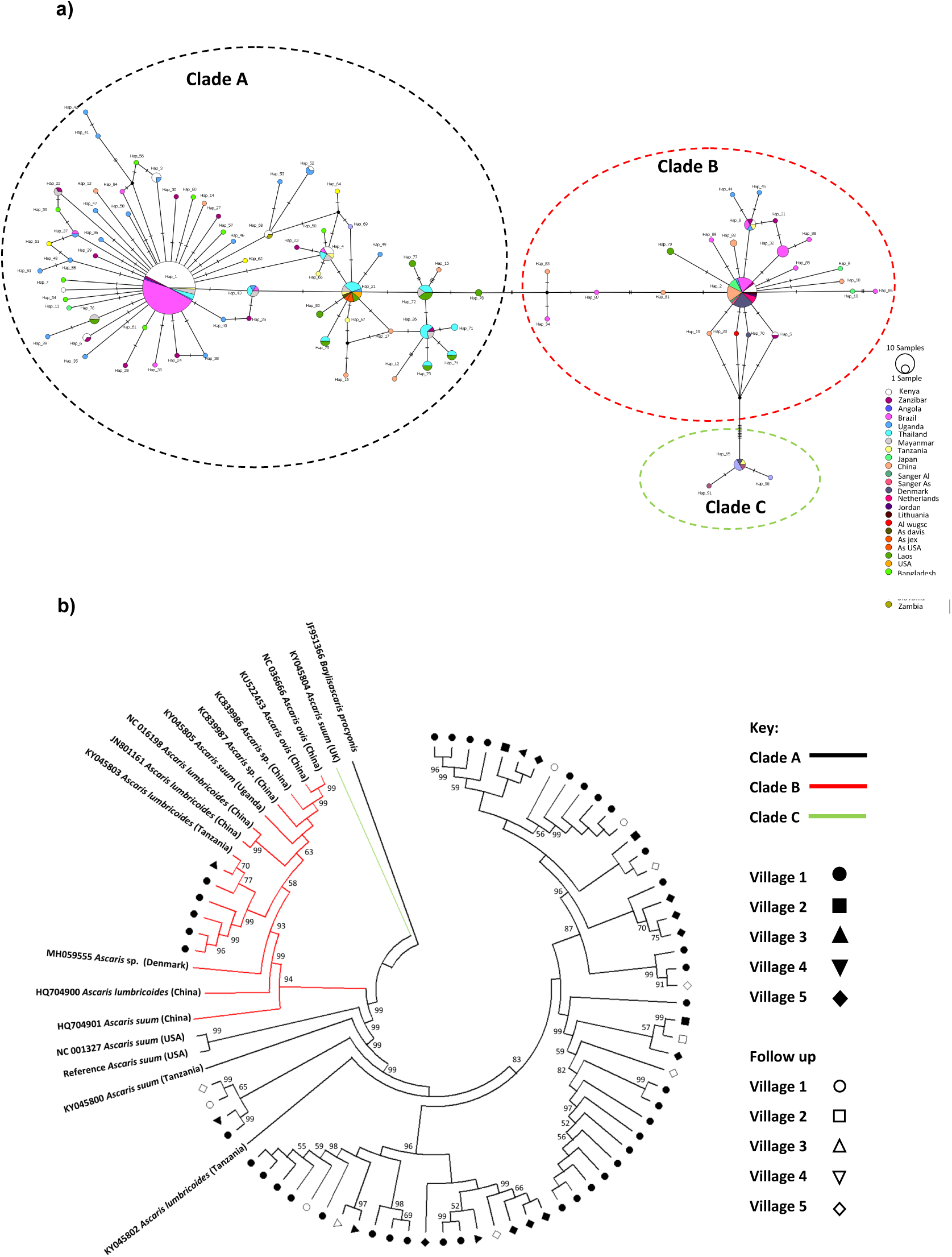
Phylogenetics of *Ascaris* spp based on mitochondrial sequences. A) Haplotype network based on the COl mitochondrial gene. Notches on the lines separating samples represent the number of nucleotide changes between the worms represented, details on the origins of haplotypes can be found in supplementary table S4; B) Maximum likelihood phylogenetic (ML) reconstruction of *Ascaris* complete mitochondrial genomes, constructed under the conditions of the GTR model and 1000 bootstrap replicates were used to provide nodal supports. The tree was constructed using all mitochondrial genomes assembled from the Kenyan isolates and all other published reference *Ascaris* mitochondrial genomes and *Baylisascaris procyonis* was used as the outgroup. The three major clades A, B and C were identified by color hue, and the majority of the Kenyan worms clustered in clade A. Each village was represented by a distinct shape and unfilled shapes represented worms sequenced from specific villages post anthelminthic treatment.

When CO1 sequences from the present study were compared against those within the *Ascaris* species complex deposited at NCBI (see Table S4 and Figure 2B)^22,23,41,42^, within clade A (which appeared to contain the majority of sequences not only from Kenya but also from other localities), 7 unique haplotypes of CO1 from Kenya were identified. These appeared to be shared not only with other haplotypes from Africa, but also with those from Brazil. In contrast, clade B haplotypes appeared to be even more cosmopolitan, with the three haplotypes from Kenya not only being shared with Zanzibar, but also with haplotypes from Brazil, Denmark, China and Japan. Despite the distinct clustering of haplotypes into the three typical *Ascaris* clades, there was very little genetic diversity among haplotypes within each of the clades, with the majority of haplotypes being separated by 1-4 nucleotide differences. There were greater levels of genetic divergence between clades; A and B were closer to each other while C was more distinct. Similar findings were seen with ND4, the most variable gene in the mitochondrial genome (Figure S2, Figure S3, supplemental text).

### Phylogenetic analyses and population structure inferred from complete mitochondrial genomes

Forty-seven SNPs were identified in the human *Ascaris* mitochondrial genomes. Approximately a quarter of these variants were in non-coding portions of the mitochondrial genome and half were synonymous (Table S1). As with the CO1 haplotype analyses, whole mitochondrial genome analysis distinguished two clades (clade A and clade B), but there were no distinct geographically specific sub-clades seen within either clade A or B (Figure 2B and Table 2). Clade C was also produced by a single published sequence which was used for comparison. In order to assess the validity of the clades A and B representing two distinct molecular taxonomic units, and thus potentially different species, Birky’s ^41^ 4X ratio was applied to provide a lineage specific perspective of potential species delimitation. The ratio failed to differentiate clades A and B as distinct species with K/θ<4 at 2.285 indicating *Ascaris* is one large population—further supporting the lack of differentiation into separate species (Table S5). Furthermore, there were no significant associations between mitochondrial sequence variations and other factors (e.g. village, household, time of worm collection, host) based on PERMANOVA (see methods and Table 2) after translating the phylogenetic tree into a distance matrix, suggesting not only a lack of differentiation into distinct species but also a potentially large interbreeding population of worms being transmitted between individuals and across villages.

**Table 2.**
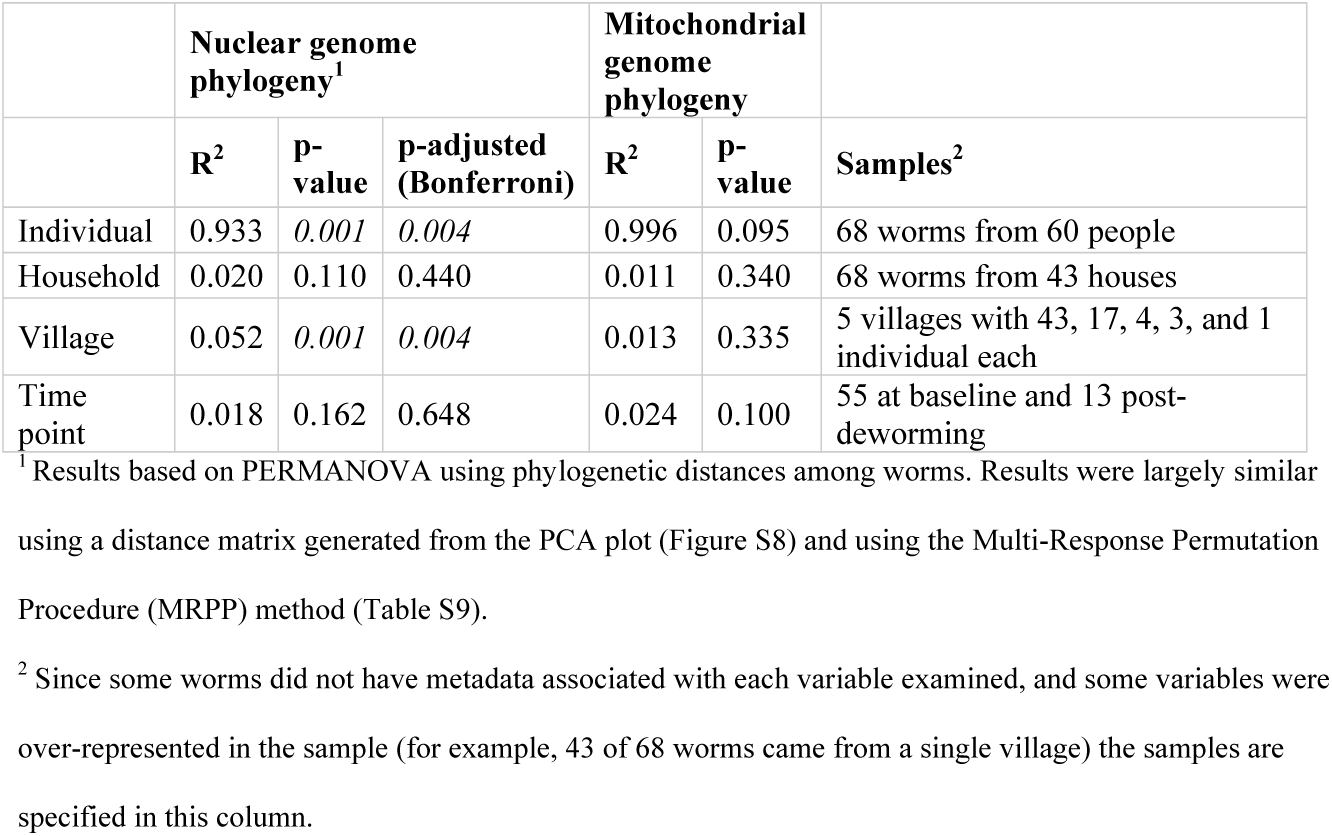
Effects of host, household, village and time point on the genetic variation of *Ascaris*.

To account for a potentially large population of interbreeding worms, analyses to detect signatures of population expansion were performed. When the global mitochondrial genome data were compared, the Tajima’s D was negative and significant (Tajima’s D −1.5691; P-value 0.028), indicating an excess of low frequency polymorphisms within the global data set suggesting population size expansion. Despite the Fu’s F not being significant it was positive (Fu’s Fs 8.5673; P-0.975) potentially indicating a deficiency in diversity as would be expected in populations that have recently undergone a bottle neck event. The same pattern was also seen in the Kenyan sequences but neither the Tajima’s D nor the Fu’s were significant. Although there does appear to be a signature of a recent population expansion event in both the global and Kenyan data, the lack of information on the mutation rates of *Ascaris* and other nematodes prevents the accurate estimate of such an event.

### Nuclear Genome Variation in the *Ascaris* population

To quantify genetic variation in the *Ascaris* worms isolated from infected Kenyans, the nuclear genomes of the 68 individual worms were analyzed to assess intraspecific population genetic diversity, heterozygosity, and ploidy. Single nucleotide polymorphisms (SNPs) and insertion/deletions (InDels) across the nuclear genomes was assessed for the first 50 largest scaffolds, that comprised 78% of the genome (see methods). Each *Ascaris* worm was sequenced to a mean coverage depth of ∼27-fold. A total of 11.15 million SNP positions were identified in the first 50 scaffolds among the *Ascaris* nuclear genomes. Approximately 25% of these variants were intergenic (Table S1). As an example, SNPs and InDels in a single *Ascaris* chromosome were plotted for two worms collected from humans in Kenya and one worm from a pig in the United States (Figure S4). The profiles and the frequency between SNPs and InDels are highly consistent within individual worms, with the ratio of InDel:SNPs frequency at ∼1:7. A comparison of the variations identified between individuals infected with worms that had either *A. lumbricoides*-like or *A. suum-*like mitochondrial genomes illustrates that most of the differences appear to be random variations, and there do not appear to be major differences between *A. lumbricoides*-like and *A. suum-*like worms. A total of 1.79 million SNPs were private, or unique to individual strains, presumably representing genetic drift. Of the remaining 9.3 million SNPs, ∼32% of these variant positions were present in less than 5 isolates indicating that the *Ascaris* genomes sequenced are ∼1% polymorphic among the major alleles circulating within the species complex.

### Population structure inferred from nuclear genomes

To investigate the evolutionary pressures that account for the high SNP diversity found among the 68 sympatric isolates, the ploidy, degree of heterozygosity (*He*) and allelic diversity was determined. Worms were disomic, with little to no evidence of aneuploidy (Figure S5). The vast majority (>98%) of SNP positions were biallelic, and each isolate had, on average, 2.3 million variant positions, of which approximately 60% were heterozygous SNPs (Table S6). SNP density was determined in 10kb windows for each worm against the reference ALV5 isolate and a patchy, mosaic pattern was resolved. SNP density was structured within the genome, with scaffolds being either SNP poor or SNP dense. For example, Algv5r020 was SNP dense whereas Algv5r019x was SNP poor. In other scaffolds, alternating SNP poor and SNP dense regions were defined within the contig, with distinct transition points, see for example the first half of Algv5b02, the last quarter of Algv5b05, or the middle of Algv5r021x (Figure 3A). In those regions where SNP density was low, the Tajima D statistic was net negative, indicating that allele frequencies within these regions were structured and more limited.

**Figure 3:**
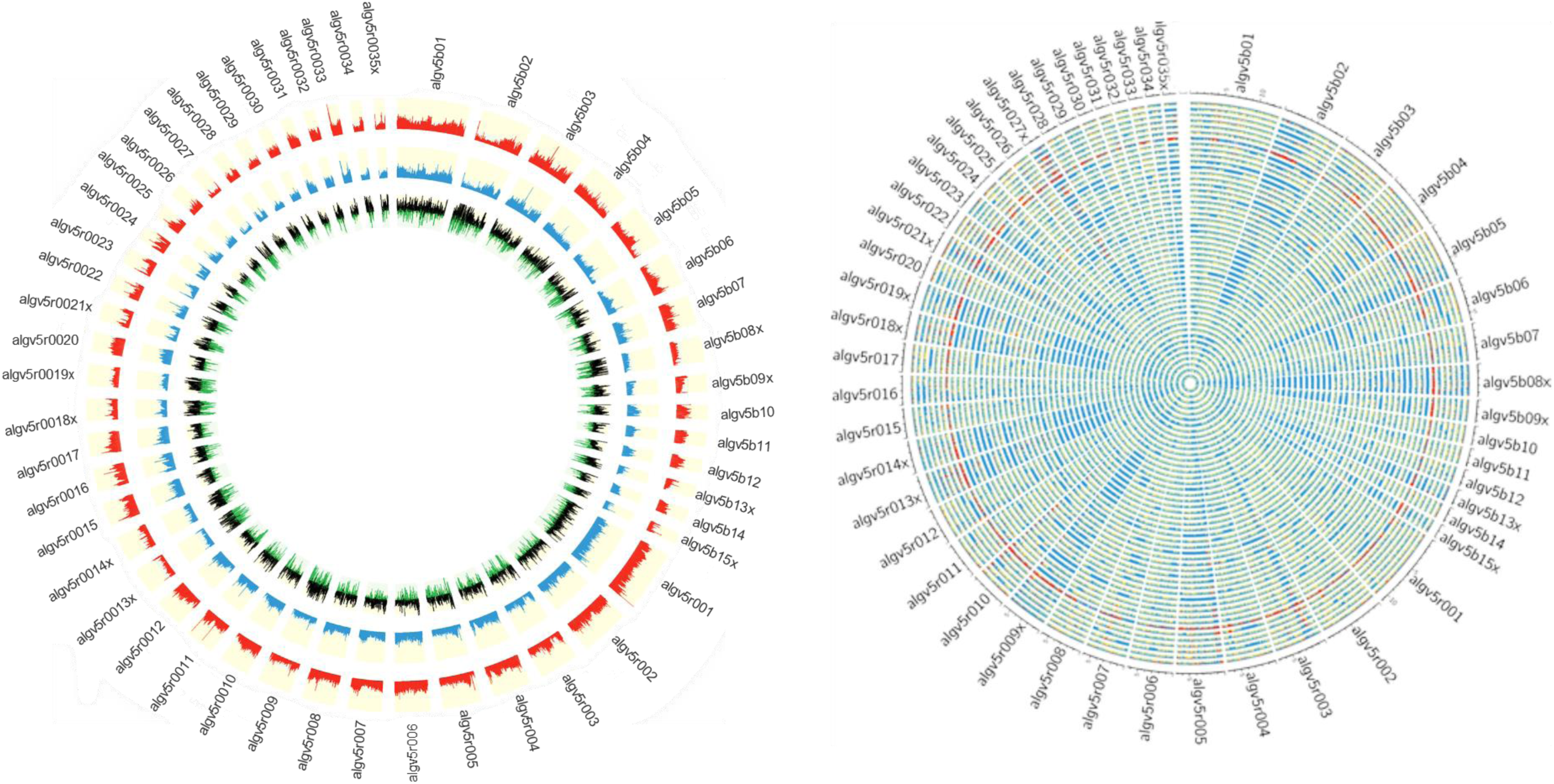
Genetic diversity of the *Ascaris* strains. A) Circos plot depicting the genetic diversity of the Ascaris strains. Outside track (red histograms) shows the total SNP diversity across the genome (first 50 largest scaffolds) in 10kb sliding windows. Blue bar plot indicates the measured degree of polymorphism (π) ^97^ within the Ascaris population in 10 kb sliding windows. The innermost track with black-green histogram plots the TajimaD ^98^ values which reflect the difference between the mean number of pairwise differences (π) and the number of segregating sites using a sliding window of 10 kb. B) The Circos-plot of the genome-wide distribution of heterozygous and homozygous SNPs in 10 kb blocks identified long stretches of homozygosity among the different strains of *Ascaris*, except 119_3, which is predominantly heterozygous throughout and was isolated from village 3. Red color = >90% of heterozygous SNPs, blue = > 90% of homozygous SNPs, yellow = 50% heterozygous, 50% homozygous SNPs. Each track represents a single strain.

Genome-wide, homozygous SNP regions were found to be unevenly distributed, with some scaffolds possessing long runs of homozygosity, see for example Algv5b02, Algv5r009x, Algv5r013x, Algv5r014x, Algv5r018x, Algv5r019x, Algv5r027x (depicted by solid blue in Figure 3B), and these regions were net negative by the Tajima D test. Conversely, heterozygous SNPs were less structured and appeared randomly distributed throughout the genome (Figure 3B). Overall, three genetic types were resolved by this analysis: in each genome there existed SNP-poor homozygous regions (colored blue) or SNP dense regions, that either possessed homozygous alternate SNPs (also colored blue) or heterozygous SNPs (colored in “red” or “yellow” blocks depending on the density of heterozygous SNPs resolved in each 10kb block: one haplotype was similar to ALV5 and the other was different). Only one isolate (119_3) was heterozygous genome-wide, and this track is depicted as “red” across all scaffolds in the Circos plot (Figure 3B).

### Population genetic structure of Kenyan *Ascaris* worm isolates

A phylogenetic tree constructed using genome wide SNPs with at least 10x coverage (11.15 million phased SNPs total) from 69 Ascaris strains, including the *A. suum* reference genome, established that the Kenyan isolates were more similar to each other than they were to the *A. suum* reference genome, which had many more private SNPs (Figure 4A). Notably, the nuclear genomes from the isolates that possessed *A. lumbricoides-*like mitochondrial genomes did not clade separately, indicating that the nuclear genomes were incongruent with the mitochondrial genomes, and likely recombinant. A co-ancestry heatmap was generated among the sympatric *Ascaris* isolates, and this analysis divided the genome into discrete segments and clustered samples along the diagonal based on the greatest number of shared ancestral blocks using the nearest neighbor algorithm from fineSTRUCTURE. The *Ascaris* genomes resolved as thirteen clusters that possessed high frequency nearest-neighbor, or shared ancestry, relationships. In contrast, the *A. suum* reference genome and strain 119_3 were anomalous, likely the result of their excess heterozygosity due in part to elevated numbers of private SNPs. Notably, 9 isolates did not coalesce into a cluster with shared ancestry. Closer examination of these strains indicated that their phased genomes possessed limited allelic diversity and were highly recombinant (Figure 4B). This genetic mosaicism was readily resolved by fluctuating intra-scaffold genealogies established using a sliding-window neighbor-joining topology that identified regions with incongruent tree topologies. See for example the trees generated at the scaffolds ALgV5b01, ALgV5b02, and ALgV5r001. Indeed, the pairwise SNP and F_ST_ estimates for these strains identified segments where SNP density was low, but F_ST_ was elevated with respect to neighboring segments (see block in ALgV5b02) and the most parsimonious explanation for these results is that recombination of a limited number of distinct alleles had occurred in the regions of increased F_ST_ (Figure 4B and 4C).

**Figure 4:**
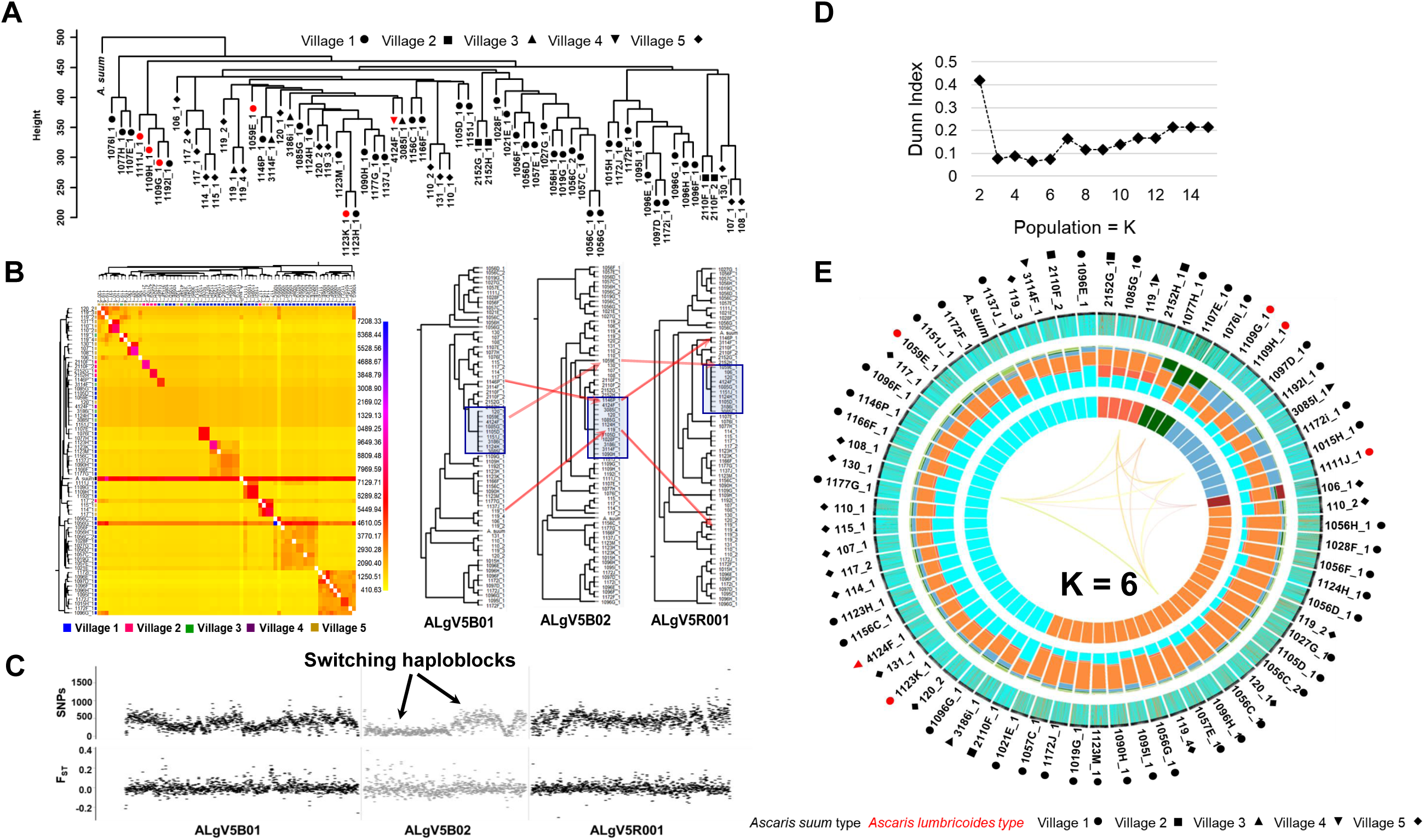
Comparative genomics and population genetic structure of *Ascaris*. A) Hierarchy phylogenetic tree of *Ascaris* strains. Phylogenetic tree was constructed with genome wide SNPs (at 10x coverage) from 68 *Ascaris* strains, including the *A. suum* reference (outgroup). Height = number of SNPs per site. Red symbol = *A. lumbricoides* mitochondrion genome. Black symbol = *A. suum* mitochondrion genome. Samples were collected from 5 different villages: Circle = village 1, square = village 2, upside triangle = village 3, downside triangle = village 4, diamond = village 5. **B)** Heatmap clustering the co-inheritance of ancestral blocks by Bayesian method using genome wide shared haplotype segments among the *Ascaris* genomes. scale = posterior coincidence probability. Hierarchical clustering and phylogenetic relationships are based on percent shared haplotype segments in scaffolds ALgV5B01, ALgV5B02, and ALgV5R001. Red arrows show examples of genetic recombination demonstrated by phylogenetic incongruence in the tree topology based on shared ancestry among blue highlighted strains (n = 13). **C)** Pairwise SNPs and F_ST_ estimates in scaffolds ALgV5B01, ALgV5B02, and ALgV5R001 indicate a switching of haplotypes (black arrow), and genetic hybridization among the blue highlighted strains (n = 13) in the phylogenetic tree depicted in figure “B”. X-axis = total SNPs/10kb in SNPs plot or F_ST_ /10kb in F_ST_ plot. **D)** Estimation of the number of ancestral populations (K) based on Dunn Index ^99^. **E)** Population genetic structure and admixture clustering analysis of the *Ascaris* genomes obtained by POPSICLE ^102^ using K=6 different color hues in the innermost concentric circle of the Circos plot. The middle concentric circle shows the relative percentage of each genetic ancestry within each genome (represented by the color hues for K = 6). The outermost concentric circle shows the genome wide local admixture profile of each strain in 10 kb sliding windows. The following geometric shapes represent villages, and the color for each shape identifies the mitochondrion genome each sample possesses: Black = *A. suum*; red = *A. lumbricoides*; Circle = village 1; square = village 2; upside triangle = village 3; downside triangle = village 4; diamond = village 5

To estimate the number of supported ancestries (K) that could be resolved in the *Ascaris* genomes sequenced, we calculated the Dunn index, which supported 3-6 ancestral populations (Figure 4D). A gradual increase in the Dunn Index after K = 6 was observed for an ancestral population size between 2 and 15 (Figure 4D and Figure S6). We next used POPSICLE to calculate the number of clades present within each 10kb sliding window. Local clades were represented with a different color and painted across the genome to resolve ancestry. The SNP diversity plots across the 68 isolates identified 3 major “parentage blocks” that were resolved as belonging to ALV5 or were genetically distinct with either both haplotypes sharing the alternate parent (homozygous alternate), or were heterozygous between the two parental haplotypes for the majority of the isolates (Figure 4E, middle Circos plot. Color hues cyan, orange, aqua).

To visualize such shared ancestry across the different *Ascaris* strains at chromosome resolution, a color hue representing a local genetic “type” present was assigned and integrated to construct haplotype blocks across each chromosome for the ancestries present. Chromosome painting based on shared ancestry revealed a striking mosaic of large haplotype blocks of different admixed color hues, consistent with limited genetic recombination between a low number of parentage haplotypes. These admixture patterns were readily visualized by shared color blocks between different strains across entire scaffolds including AlgV5R019X (Figure 5A) and AlgV5R027X (Figure 5B). In low complexity regions such as the left portion of contig ALgV5R019X, only three major haplotypes were resolved (Figure 5A). Strikingly, within each of the 6 clades resolved, all strains showed a limited, mosaic fingerprint of introgressed sequence blocks indicating that recombination has shaped the population genetic structure among the *Ascaris* isolates sequenced. Evidence for both segregation and recombination were evident. For example, isolates 1107E_1 and 2110F_2 shared the same chromosome at ALgV5R019X, but entirely different chromosomes at ALgV5R027X, whereas isolates 107_1, 108_1 and 2110F_2 were identical except at the subtelomeric end of ALgV5R19X. In this region two admixture blocks were resolved; 107_1 and 2110F_2 remained similar to each other but 108_1 now possessed a sequence block that was shared with isolate 119_3. This extensive chimeric pattern in chromosome painting also closely resembled the genome-wide hierarchy tree (Figure 5A). The data support a model in which the isolates are genetic recombinants between *A. suum* and *A. lumbricoides* that are predominantly inbreeding.

**Figure 5:**
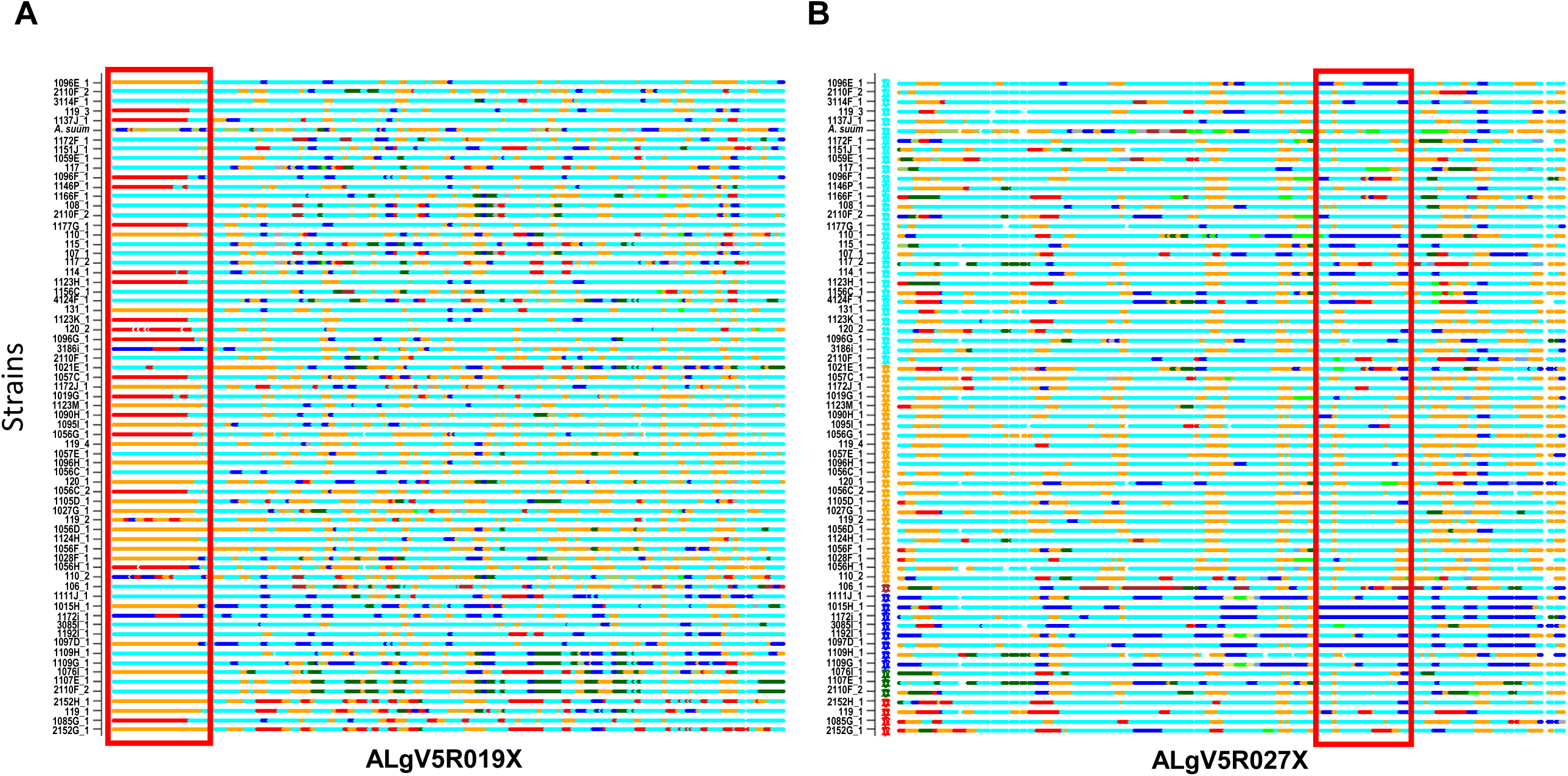
Local admixture clustering and genome wide analysis of inheritance of haploblocks of *Ascaris* obtained by POPSICLE ^104^. Based on ancestral population K= 6. X-axis = strains. Red highlighted box indicates the introgression of large haplotype blocks of defined parentage among the different strains of *Ascaris* in scaffolds ALgV5R019X (A) and ALgV5R027X (B). Many examples exist whereby strains that are in linkage disequilibrium at ALgV5R019X possess different haplotypes in ALgV5R027X (for example 1107E_1 vs. 2110F_2) indicating both segregation as well as recombination in the evolution of the isolates. The local admixture patterns reveal extensive genetic hybridization among different strains of *Ascaris*. Color assignment is depicted based on Figure 6E.

### Geographic and demographic correlates of genetic similarity

To examine clustering of worms in similar human hosts, we statistically compared genetic variation within groups (such as within a village) versus between groups (such as between villages). We found significant genetic separation between worms in different villages (using Adonis vegan in R), but not between worms from different countries (Table 2). This suggests genetic diversity is present in the population of *Ascaris* in these Kenyan villages, which is similar to the diversity of populations of *Ascaris* around the world. It also suggests that a high proportion of *Ascaris* transmission may occur within villages in this Kenyan setting. There was no evidence from this analysis that the 13 worms collected three months after albendazole treatment were any different than the worms collected prior to albendazole treatment (Table 2).

To expand on our observations, that genetically similar worms are found around the world, but that similar worms cluster within a village based on our nuclear SNPs data, we plotted genetic distances against geographic distances. Surprisingly, we found no significant correlations between genetic and geographic distance, neither across all five studied villages nor within the two most heavily parasitized villages (Figure S7 and supplemental text).

## Discussion

In this study, we generated a high-quality reference genome from a single worm presumed to be human *A. lumbricoides*. Our comparative phylogenomic analyses of this new *Ascaris spp.* genome against existing draft genomes of *A. lumbricoides* and *A. suum* suggest that *A. suum* and *A. lumbricoides* form a genetic complex that is capable of interbreeding that has apparently undergone a recent worldwide, multi-species *Ascaris* population expansion.

Our phylogenetic analysis on the complete mitochondrial genomes (from 68 worms collected from human hosts in Kenya and other available sequences) suggests that the worms collected in Kenya mirrors the separation into clade A (worms from pigs in non-endemic regions and humans in endemic regions) and clade B (worms from humans and pigs from endemic and non-endemic regions) described elsewhere ^22^. It is likely that worms in both of these clades are being transmitted from human to human, as pig husbandry is rare in this area of Kenya. Patterns may differ by locality, and it is possible that some of the pig-associated (*A. suum*-like) worms circulating in this human population in Kenya were acquired, perhaps generations ago, by humans who lived in closer proximity to pigs. It is also possible that these worms were acquired from non-human primates ^42^, or some other *Ascaris* host, rather than from pigs.

However, the SNPs across the whole nuclear *Ascaris* genome provide significantly greater power in understanding *Ascaris* speciation Importantly, our nuclear genome SNP analysis suggest that the 68 Kenyan *Ascaris* are distributed across multiple clades in a phylogeny based on the nuclear genomes. Overall, data from our study and other studies are consistent with a pattern where hybrid genotypes in *Ascaris* populations were observed ^11,22,43^ Our study represents one of the most detailed accounts of mito-nuclear discordance in nematodes echoing patterns seen in another human nematode *Onchocerca volvulus* ^44^. The data in our current study shows the occurrence of distinct mitochondrial lineages that could be evidence of early stages of species differentiation. The admixture seen within the nuclear genome, however, appears to disrupt the establishment of defined molecular speciation barriers between the different *Ascaris* lineages. Such patterns have been recorded in other parasites not only in *O. volvulus* ^44^, but also in the parasitic blood fluke *Schistosoma* ^45^ and the protist *Leishmania* ^46^. Each of the studies have implicated definitive hosts in the movement of parasites between otherwise isolated populations allowing interbreeding to take place. It is most likely the historical movement of humans and their domesticated livestock that has mediated the transport of *Ascaris* between localities, allowing for extensive interbreeding within the nuclear genomes and the discordance observed between the mitochondrial and nuclear genomes in our study.

At a more local scale, the insights into the human transmission dynamics of *Ascaris* showing clustering both within an individual and in villages suggest that villages are appropriate units for interventions and that people are infected with multiple eggs from a single source. These findings are in line with clustering at the village level found in Guatemala ^47^ and at the sub-village level in Nepal ^48^, but not in line with the lack of small-scale geographical structuring found in Denmark, Zanzibar and Uganda ^49–51^. Differences could be a result of different patterns in human and livestock movement ^20^.

Although the current genome is, by far, the most continuous assembly for *Ascaris*, it is not a full chromosome assembly due largely to repetitive sequences, in particular 120 bp tandem repeat clusters and long stretches of subtelomeric repeats. Thus, it is possible that mis-assembly in some scaffolds has increased the frequency of mosaicism detected. It is for this reason that the comparative analyses on the nuclear genome was restricted to the largest 50 scaffolds, most of which are at chromosomal resolution, with only minor localized variation due to the repeat clusters. In these high confident scaffolds, large haplotype blocks possessing either *A. suum, A. lumbricoides* or both parental haplotypes (heterozygous) were readily resolved indicating that the genetic mosaicism observed could not be solely attributed to genome mis-assembly. Ultimately, future studies using ultralong PacBio and/or Nanopore sequencing combined with chromosome conformation capture (Hi-C) techniques will improve the genome to full chromosome assembly to more accurately resolve the true extent to which recombination has impacted the population genetic structure of the *Ascaris* species genetic complex.

The finding that *A. suum* and *A. lumbricoides* form a genetic complex has important public health implications. Reduced treatment efficacy is not currently a common issue in *Ascaris* infections among humans or pigs ^52–54^, though low efficacy of benzimidazoles is an issue for *Trichuris trichiura* in humans ^55–57^ and various intestinal nematodes of veterinary importance ^58–60^. Extensive albendazole use in either human or pig populations could lead to resistance in both populations, if cross-species infections are common and produce fertile offspring. This study suggests that research and public health interventions targeting *A. lumbricoides* and *A. suum* should be more closely integrated, and that extensive work done by the veterinary research community may be highly relevant to mass deworming campaigns that seek to improve human health.

The similarity between *Ascaris* from different countries and from different vertebrate hosts suggests that *Ascaris* infection has spread rapidly around the world, leaving little time for it to differentiate. Taken together, these finding have very important implications for parasite control and elimination efforts that only focus on mass deworming of humans for *Ascaris*. The ability of pig-associated worms to become endemic in human populations indicates that a one-health approach may be necessary for the control of *Ascaris*.

## Online Methods

### Worm collection

Worms were expelled as part of a larger study in rural western Kenya described previously ^61,62^. Worms collected from study participants in five villages (map shown in ^61^) following treatment with 400 mg albendazole were isolated, washed, labelled and stored frozen (−15 C). They were transported from Bungoma to Kisumu, where they were subsequently stored at the KEMRI-CDC offices until they were shipped to the NIH (Bethesda, MD, USA) on dry ice.

### DNA extraction and sequencing

A modified DNA extraction method was developed (see Supplemental Methods and Table S3). For the five germline samples, DNA was extracted from the uterus, oviduct or ovary of the worms. For the remaining samples, DNA was extracted from somatic tissue: the body wall or the intestine. Paired-End Genome Libraries – Sixty-eight *A. lumbricoides* DNA samples were sequenced using Illumina HiSeq 2500 (www.illumina.com) short-read paired-end sequencing. DNA was quantified by UV Spec and Picogreen. A 100 ng of DNA based on picogreen quantification was used as template for NGS library preparation using the TruSeq Nano DNA Sample library prep kit without modification. Primer-dimers in the libraries were removed by additional AMPure beads purification. Sequencing was performed to obtain a minimum genomic depth of 20X coverage for each sample.

Mate-Pair Genome Libraries – Two samples were selected for mate-pair sequencing, based on the quality of the DNA preparation. Three independent DNA isolations (corresponding to what region of the worm or what is the sample for DNA isolation) from specimen “119_2.3” were combined to obtain one μg DNA input. The mate-pair libraries were generated using the Nextera Mate Pair Library Prep Kit, following the gel-free method with the only modification that M-270 Streptavidin binding beads were used instead of M-280 beads. The libraries were amplified for 15 cycles given the low DNA input going into the circularization phase. The mate-pair fragment size averaged 6 kb with a range of 2-10 kb fragments.

### Assembly and annotation of A. lumbricoides reference genome

The *A. lumbricoides* germline genome assembly was constructed using the *A. suum* genome as a reference. Briefly, sequencing reads from a single *A. lumbricoides* worm (libraries #8457, #8458 and #8778) were mapped to the *A. suum* germline genome assembly ^28^ using BWA^63^ to generate BAM and MPILEUP alignment files. The MPILEUP files were processed with a PERL script that replaced all variation sites in the reference genome with the highest allele frequencies in the *A. lumbricoides* sample. *A. suum* genomic regions that represent < 5X of *A. lumbricoides* reads coverage were excluded from the assembly. We further polished the genome with additional Illumina sequencing reads using Pilon and its default parameters ^64^. The *A. lumbricoides* genome was annotated using the gene models built for *A. suum*, using the annotation transfer tool RATT ^65^. The protein coding regions were defined using TransDecoder (https://github.com/TransDecoder/TransDecoder/wiki). To evaluate the gene expression across all stages, we utilized previous RNAseq data from the developmental stages ^27,28^, re-mapped the SRA from adult males, females, L3 and L4 stages ^25^ to the current gene models, and quantified the expression using tophat and cufflinks. The re-mapped reads, analyzed by JMP Genomics (SAS) across all the stages and based on the principal component analyses (Figure 1B), were grouped as adult male, adult female, L1, L2, L3 (egg L3, liver L3 and lung L3), L4, carcass, muscle, intestine, embryonic (zygote1, zygote2, zygote3, zygote4, 24h, 46h, 64h, 96h, 5d, 7d), ovaries (female mitotic region, female early pachytene, female late pachytene, female diplotene and oocyte) and testis (male mitotic region, spermatogenesis, post meiotic region, seminal vesicles and spermatids). Proteome and comparative genomics analyses were done using an in-house pipeline ^66^. Automated annotation of proteins was done as described earlier ^31^ and based on a vocabulary of nearly 290 words found in matches to various databases, including Swissprot, Gene Ontology, KOG, Pfam, and SMART, Refseq-invertebrates and a subset of the GenBank sequences containing nematode protein sequences, as well as the presence or absence of signal peptides and transmembrane domains. Signal peptide, SecretomeP, transmembrane domains, furin cleavage sites, and mucin-type glycosylation were determined with software from the Center for Biological Sequence Analysis (Technical University of Denmark, Lyngby, Denmark) ^67–69^. Classification of kinases was done by Kinannote ^70^. Interproscan ^71^ analyses were done using the standalone version 5.34. Allergenicity of proteins were predicted by Allerdictor ^72^, FuzzyApp ^73^ and AllerTOP ^74^. Genes that had blast scores <30% of max possible score (self-blast) in other non-Ascaris nematodes with an e-value greater than 1E-05 were considered as ‘unique’. The orthologues of predicted proteome of ALV5 across the publicly available nematode genomes (*Ancylostoma caninum* ^32^, *Ancylostoma ceylanicum* ^32,75^, *Ancylostoma duodenale* ^32^, *Ascaris lumbricoides*^32^, *Ascaris suum* ^25,27,28^, *Brugia malayi* ^76^, *Caenorhabditis elegans* ^77^, *Dirofilaria immitis* ^78^, *Loa loa* ^79,80^, *Necator americanus* ^81^, *Onchocerca volvulus* ^31^, *Strongyloides ratti* ^82^, *Strongyloides stercoralis* ^83^, *Toxocara canis* ^32,84^, *Trichinella spiralis* ^85,86^, *Trichuris trichiura* ^87^, *Wuchereria bancrofti* ^32,88^) were analyzed using OrthoFinder ^89^. The estimated phylogenetic tree generated was graphed using FigTree v1.4.

Further manual annotation was done as required. The data were mapped into a hyperlinked Excel spreadsheet as previously described ^90^, available in Table S2.

### Read mapping and SNP analysis for whole genome sequences

The Illumina paired end sequence reads of the 68 *Ascaris* whole genomes were trimmed by removing any adapter sequences with CutAdapt v1.12 ^91^, then low quality sequences were filtered and trimmed using the FASTX Toolkit (http://hannonlab.cshl.edu/fastx_toolkit/). Remaining reads were then ref-mapped to the *A. lumbricoides* genome ALV5 reference genome (described in this paper) using either Bowtie2 v2.2.9 ^92^, with very sensitive, no-discordant, and no-mixed settings or using the Burrows-Wheeler Aligner (BWA, v0.7.9) ^63^ mem in default parameters and then converted into a bam file for sorted with SAMtools ^93^. Sorted reads were soft-clipped and marked-duplicated using Picard-1.8.4 (http://broadinstitute.github.io/picard). Single nucleotide polymorphisms (SNPs) were obtained using SAMtools ^93^ and BCFtools ^94^ using the mpileup function and –ploidyfile features and taking chromosomal ploidies into account. SNPs were also determined using Genome Analysis Toolkit (GATK) ^95^. SNPs were called by GATK Haplotype Caller with a read coverage ≥ 10x, a Phredscaled SNP quality of ≥30. Mapping statistics were generated in Perl and Awk.

### Ploidy determination

The ploidy of each isolate was calculated using AGELESS software (http://ageless.sourceforge.net/) by dividing the chromosomes into 10kb sliding windows and averaging the coverage within each window. The windows with zero coverage were not included in any further analyses due to sequencing noise or repeat regions ^96^.

### Genetic diversity

SNPs, pi ^97^, TajimaD ^98^, and F_ST_ ^99^ values were calculated using VCFtools ^100^ in 10 kb sliding windows and plotted using either Circos ^101^ or ggbio (http://bioconductor.org/packages/release/bioc/html/ggbio.html) and VariantAnnotation (http://bioconductor.org/packages/release/bioc/html/VariantAnnotation.html) R packages (v. 3.1.0, URL http://www.R-project.org). The proportions of heterozygous and homozygous SNPs were estimated in 10kb sliding windows using custom Java scripts to generate histogram plots in Circos ^101^. Red and blue colors indicate the presence of 90% or more heterozygous and homozygous SNPs respectively whereas yellow color was assigned otherwise.

### Co-ancestry heatmap

The SNP data (VCF file) was first phased accurately to estimate the haplotypes using SHAPEIT ^102^ after keeping only biallelic SNPs and loci with less than 80% missing data. Co-ancestry heatmaps were generated using the linkage model of ChromoPainter ^103^ and fineSTRUCTURE (http://www.paintmychromosomes.com) based on the genome-wide phased haplotype data. For fineSTRUCTURE (version 0.02) ^103^, both the burn-in and Markov Chain Monte Carlo (MCMC) after the burn-in were run for 1000 iterations with default settings. Inference was performed twice at the same parameter values.

### Population genetic structure

Population genetic structure was constructed using POPSICLE ^104^ by comparing strains against the reference sequence ALV5 in 10 kb sliding windows with the number of cluster K=1 to 15 and then use the Dunn index ^99^ to calculate the optimal number of clusters. After calculating the optimal number of clusters, POPSICLE assigned each block to the existing or new clades depending on population structure of each strain and the ancestral state of each block followed by painting in Circos plot ^101^ with color assignment based on number of clusters.

### Construction of phylogenetic trees

In order to determine the phylogenetic relationship between samples, we selected 19005 base positions where variants were detected in a representative sample vs the reference (ALV5), and where each sample had at least 20x coverage for each locus. Using this list, the base calls for each sample were pooled together to generate a single multi sequence fasta file.

Next, both maximum likelihood (ML) trees and bootstrap (BS) trees were generated with a final “best” tree generated from the best scoring ML and BS trees using RAxML v8.2.10 ^105^. The tree was visualized in FigTree v1.4.3 (http://tree.bio.ed.ac.uk/software/figtree/).

### Permutational Multivariate Analysis of nuclear phylogeny

Similarity within and between worms from different villages, households, people and time-points was analyzed based on the distance matrix of the patristic distances from the phylogenetic tree described above, using permutational multivariate analysis of variance (Adonis Vegan in R). The distance matrix underlying the phylogenetic tree was analyzed in order to measure the significance and contribution of different factors to variance between samples. Each factor (village, household, host, time-point, human age group and worm age/size group) was analyzed both separately and sequentially. The sequence chosen was ordered based on significance of each factor when tested individually. Since multiple groupings were considered using the same dataset, multiple comparison corrections were applied. Sample sizes and descriptions of each group are shown in Table 2. Worms with recorded lengths were split into three groups of equal number. This was in order to test whether larger worms (using worm length as a very approximate proxy for age, where older worms might have survived at least one round of ALB treatment) were any different from the newly acquired younger worms. Similar methods were used to analyze the mitochondrial phylogeny along the same groupings.

### Mitochondrial genome assembly

We assembled mitochondrial genomes using a *de novo* approach from 68 individual *Ascaris* genomes. For each individual, the *Ascaris* mitochondrial reads in the total DNA sequencing were identified by mapping the *Ascaris* reads to the *A. suum* reference mitochondrial genome (GenBank accession: NC_001327). Adaptor sequences were trimmed prior to *de novo* assembly. To reduce the complexity of the *de novo* assembly, we randomly sampled 1,000x reads from each individual (the use of higher read coverage often resulted in fragmented scaffolds) and assembled these reads using the SPAdes assembler ^106^ with continuous k-mer extension from K=21 to the maximum k-mer allowed (average extended k-mer size = 91). The assembled scaffolds were corrected with the built-in tool in SPAdes to reduce potential assembly artifacts. Next, the assembled scaffolds were aligned to the *A. suum* mitochondrial reference genome using BLAST, the order of the scaffolds was adjusted, and they were joined into a single scaffold. Finally, the gaps in the scaffold were filled using GapFiller ^107^ using mitochondrial reads from the same individual to generate a complete mitochondrial genome. Using the same method, we also *de novo* assembled another five *A. suum or A. lumbricoides* mitochondrion genomes from previous studies (see Table S7).

### Analysis of mitochondrial genomes

In order to assess overall evolutionary relationships across the complete mitochondrial genomes, we aligned the genomes using Clustal W and phylogenetic trees constructed using RaxML under the conditions of the general time reversible model (GTR) as described above for the whole genome SNP alignment. Subsequent tree files were formatted in FigTree and MEGA v7. The variation in nucleotide diversity across the mitochondrial genome was measured using sliding window analyses, with a window of 300 bp and a step of 50bp, using DNAsp v6 ^108^. In order to assess the validity of potential species groupings in the ML phylogenetic tree the Birky ^41^ X4 ratio was applied to the alignment of the complete mitochondrial genomes including both samples from Kenya and published mitochondrial reference genomes from Tanzania, Uganda, China, USA, Denmark and the UK. The X4 ratio method of species delimitation compares the ratio of mean pairwise differences between two distinct clades (K) and the mean pairwise differences within each of the clades being compared (θ). It is considered that if K/θ>4 this is indicative of the two clades representing two distinct species. Owing to the fact that two clades are being compared there will be two separate values of θ, as per recommendations of Birky ^41^, the larger θ value is used to perform the final ratio calculation as this will provide a more conservative result which ultimately will be less likely to provide a false positive result.

Due to the extensive use of mitochondrial genome data in population genetic analyses of *Ascaris*, several analyses were performed to identify the effect of any population level processes that may be affecting the diversity of the parasites within Kenya. Initially, diversity indices were calculated for each of the genes within the mitochondrial genome across the entire Kenyan data set as well as considering the mitochondrial genome as a whole. In order to account for the diversity within the genic regions, we removed non-coding and tRNA sequences for these analyses. To provide a genealogical perspective of population structure of the Kenya *Ascaris* isolates, we constructed the most parsimonious haplotype network based on the protein coding sequences using the TCS algorithm as implemented in PopArt ^109^. Further population genetic analyses were also performed to detect the occurrence of selection on the protein coding genes of the mitochondrial genome and if there were any major departures from neutrality. Standard dN/dS ratios were performed to identify the presence of positive selection where both measures equate to 1 = neutral, >1 = positive selection, <1 = purifying selection. Both Tajima’s D and Fu’s *Fs* were calculated to identify any substantial departure from neutrality which could be indicative of population expansion events (Table S8). All described analyses were performed using DNAsp6 ^108^. As both CO1 and ND4 have been used in the past for epidemiological studies, single gene phylogenies were also constructed as described previously for comparison against the whole mitochondrial genome phylogeny (Figure S3 and supplementary methods).

Owing to the extensive use of the CO1 for epidemiological studies the gene was extracted from the complete mitochondrial genomes of Kenya and compared to all other available *Ascaris lumbricoides* and *Ascaris suum* CO1 sequences housed by NCBI representing populations from across the globe. Haplotype network analyses was performed to produce the parsimonious network using TCS as implemented through PopArt ^109^. This provided a genealogical perspective of population structure and allowed genetic connectivity between the Kenyan samples and other geographical isolates to be assessed.

## Acknowledgements

We would like to thank the school children, schoolteachers, and Bungoma administrators for their support. We would like to extend special thanks to all the members of the study team: Bungoma County Hospital, Siangwe, Siaka, Sang’alo, Nasimbo and Ranje village administrators and Community Health Workers. Particular thanks to Dr. Charles S. Mwandawiro, Prof. Sammy Njenga, and Dr. Jimmy H. Kihara (KEMRI), and Dr Simon J. Brooker (BMGF) for making the fieldwork possible in Kenya, and for their invaluable scientific and logistical advice.

## Data availability

Data are available under the National Center for Biological Information (NCBI) BioProject numbers; PRJNA511012 and SRA submission SUB4949491 for sequencing data, and PRNJA515325 for the genomic assembly.

Links to all genome assemblies are available at: <u>All</u>, <u>de novo</u>, <u>semi-de novo (V1)</u>, <u>V2</u>, <u>V3</u>, <u>V4</u>, <u>V5</u> and <u>mitochondrial</u>.

## Funding

This work was supported in part by the Division of Intramural Research (DIR) of the National Institute of Allergy and Infectious Diseases, NIH and NIH grants to R.E.D. (AI114054) and J.W. (AI125869). Fieldwork was supported by a grant from the Bill and Melinda Gates Foundation to the London Centre for Neglected Tropical Disease Research and the KEMRI Wellcome Trust. The funders had no role in study design, data collection and analysis, decision to publish, or preparation of the manuscript.

## Authors’ contributions

AVE, RMA, RED, JW, TBN conceived the study; SG, ED, JW, ED, SFP, RED performed sequencing and genome analyses; SPL, JPW, AK, MEG performed population biology and evolutionary biology expertise and analyses; AVE, SK, RGO, RMA oversaw the collection of material in Kenya, SB, SG, ED, JW provided protein/gene content analyses, AVE, SPL, SB, AK, ED MED, RED, JW, TBN wrote the manuscript.

## Ethics approval

This study was approved by the Ethics Review Committee of the Kenya Medical Research Institute (Scientific Steering Committee protocol number 2688) and the Imperial College Research Ethics Committee (ICREC_ 13_1_15). Informed written consent was obtained from all adults and parents or guardians of each child. Minor assent was obtained from all children aged 12–17. Anyone found to be infected with any STH was treated with 400 mg ALB during each phase of the study, and all previously-untreated village residents were offered ALB at the end of each study phase.

## Consent for publication

Individuals consented to the publication of their results, without any patient identifying information.

## Competing interests

The authors declare that they have no competing interests. RMA was a Non-Executive Director of GlaxoSmithKline (GSK) during the period of worm collection in Kenya. GSK played no role in the funding of this research or this publication.

## Supplemental Figure Legends

**Figure S1.**
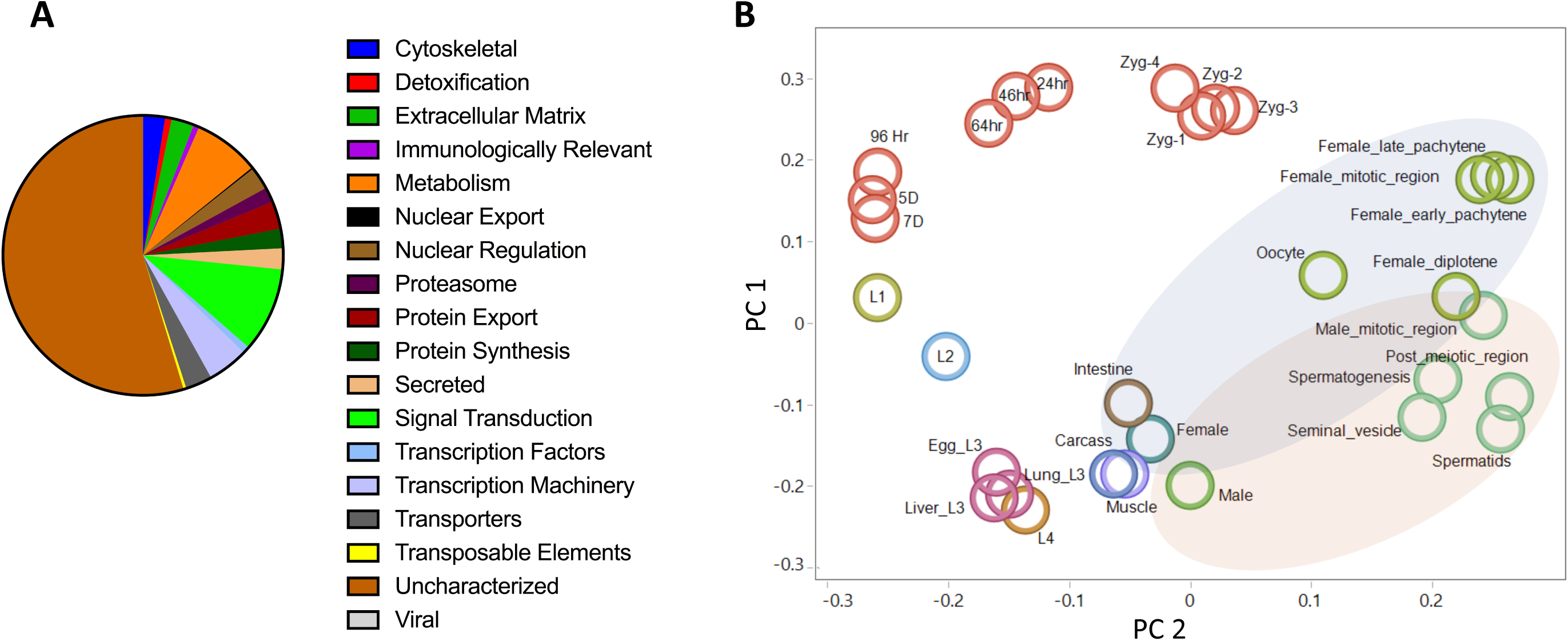
Predicted proteome and stage-specific transcriptomes of Ascaris. A) Functional classification of the predicted proteome of *A. lumbricoides* (an improved proteome of *Ascaris* spp.) with the majority of proteins being unknown/uncharacterized. B) 2-dimensional principal component analysis plot illustrating the similarities in transcription profiles between the major stages (Fig 1B) and the developmental stages.

**Figure S2.**
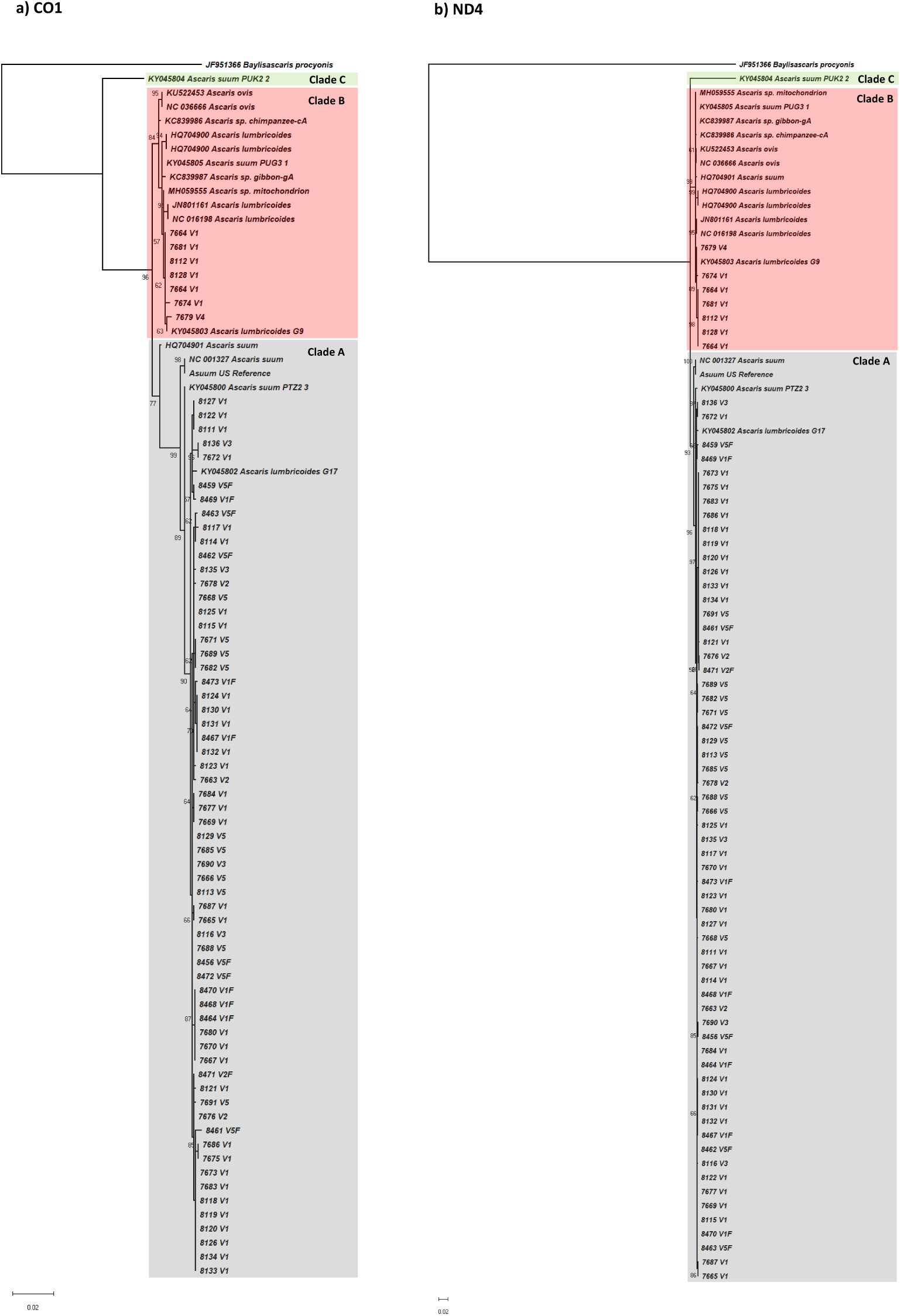
Phylogenetic trees based on CO1 and ND4. Maximum likelihood phylogenetic analyses of the A) CO1 and B) ND4 genes using RaxML under the conditions of the GTR model with nodal support values generated through 1000 bootstrap replicates. The trees were generated using complete sequences of the genes extracted from the 68 Kenyan *Ascaris* mitochondrial genomes generated in this study and other published reference genomes.

**Figure S3.**
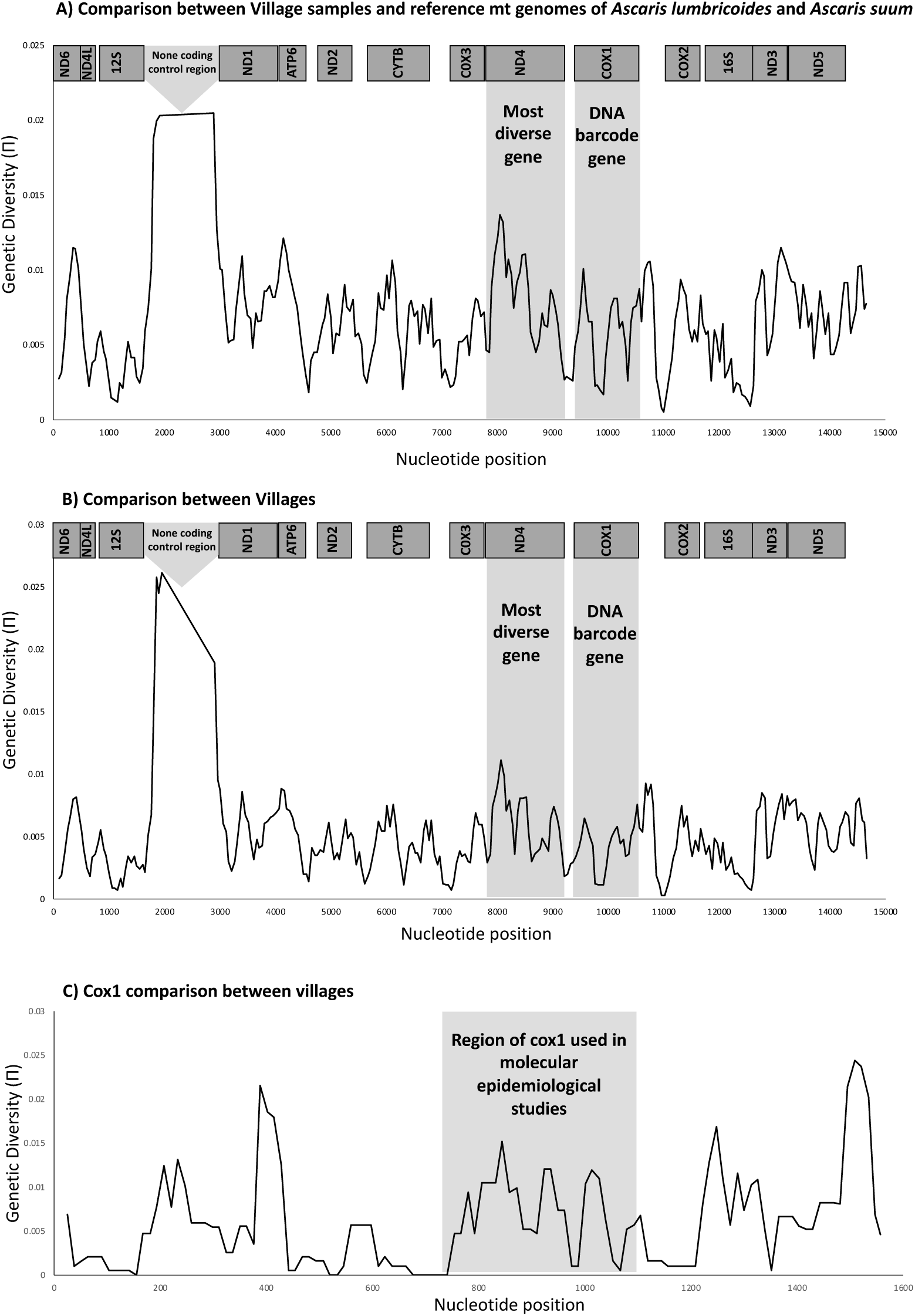
Sliding window analyses. A) Comparison between Kenyan samples and reference mitochondrial genomes of *Ascaris lumbricoides* and *Ascaris suum*, B) Comparison between villages, C) CO1 comparison between villages.

**Figure S4.**
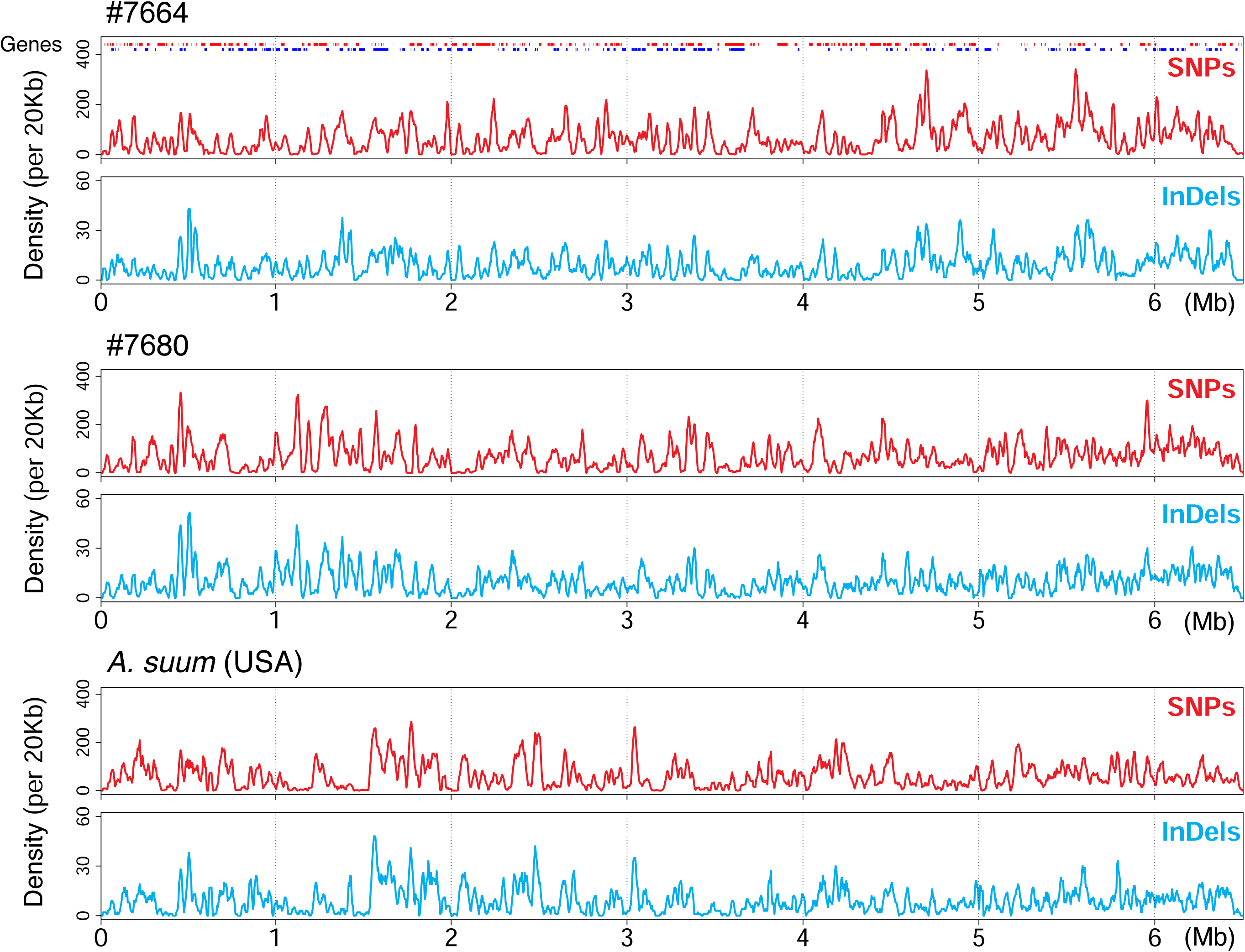
*Ascaris* SNPs and insertion/deletions (InDels) maps. An assembled 6.5 Mb *Ascaris lumbricoides* chromosome fragment (ALgV5R006), with the frequency of identified SNPs and InDels plotted for one representative *A. lumbricoides-like* worm from this study (#7664) and one *A. suum-like* worm (#7680). Genes are shown on the top of the plot, with red and blue indicating genes transcribed from forward and reverse strands, respectively. The y-axis shows the frequency of SNPs and InDels for a 20-kb window size (with a 4-kb sliding window in x-axis). Note the profiles and the frequency between SNPs and InDels are highly consistent within individual worms.

**Figure S5.**
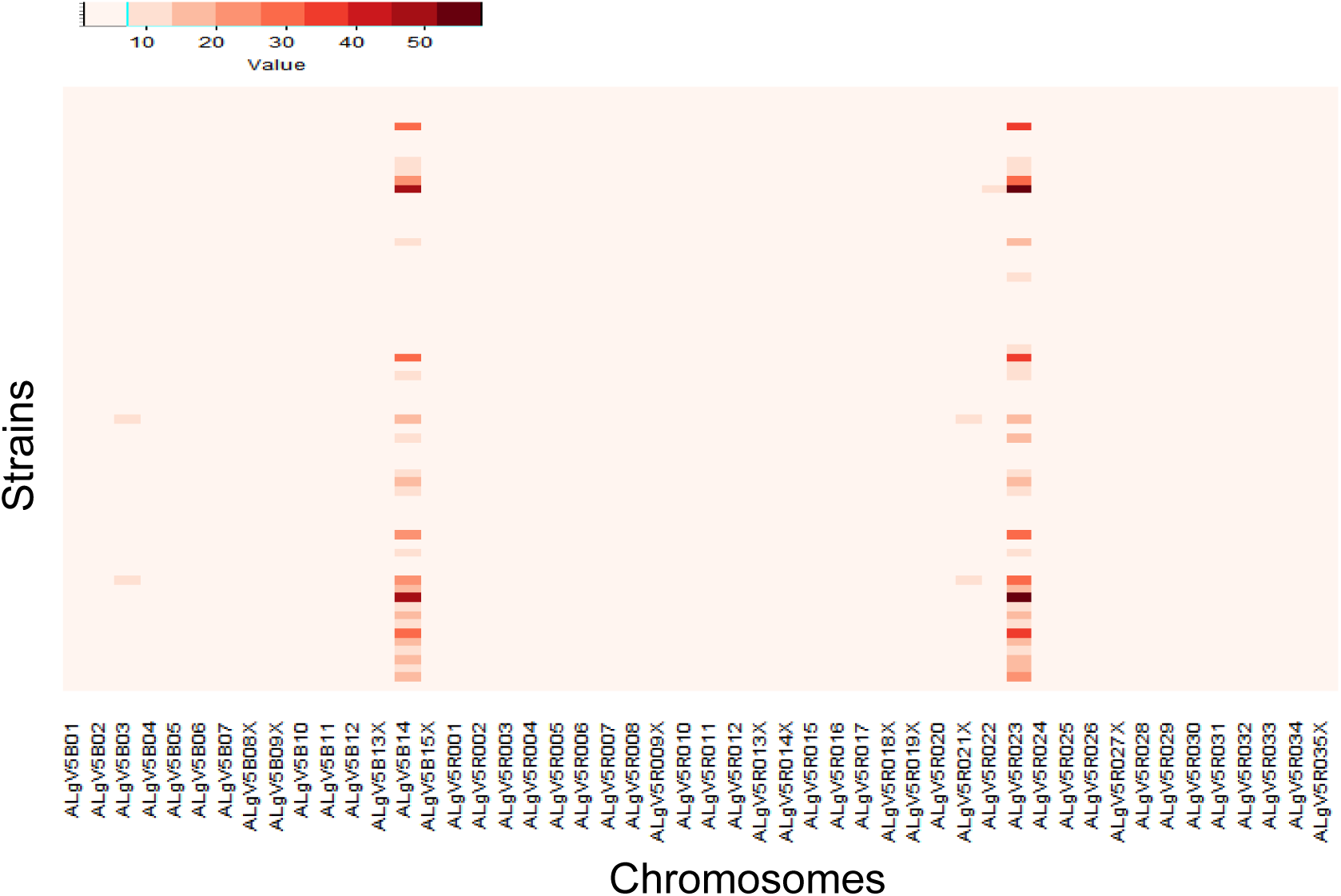
Somy analysis of the *Ascaris* isolates. The ploidy of the *Ascaris* strains are represented in a heatmap. Ploidy was calculated by averaging the count of aligned reads in 10 kb sliding windows across the genome after reference mapping against ALV5. The ploidy data suggest that *Ascaris* is completely diploid (close to 2n), except at two scaffolds ALgB5B14 and ALgv5RO23, where the majority of strains show elevated ploidy. X-axis shows the first 50 largest scaffolds involved in this study and the y-axis shows the strains (ordered by code number 1-68).

**Figure S6.**
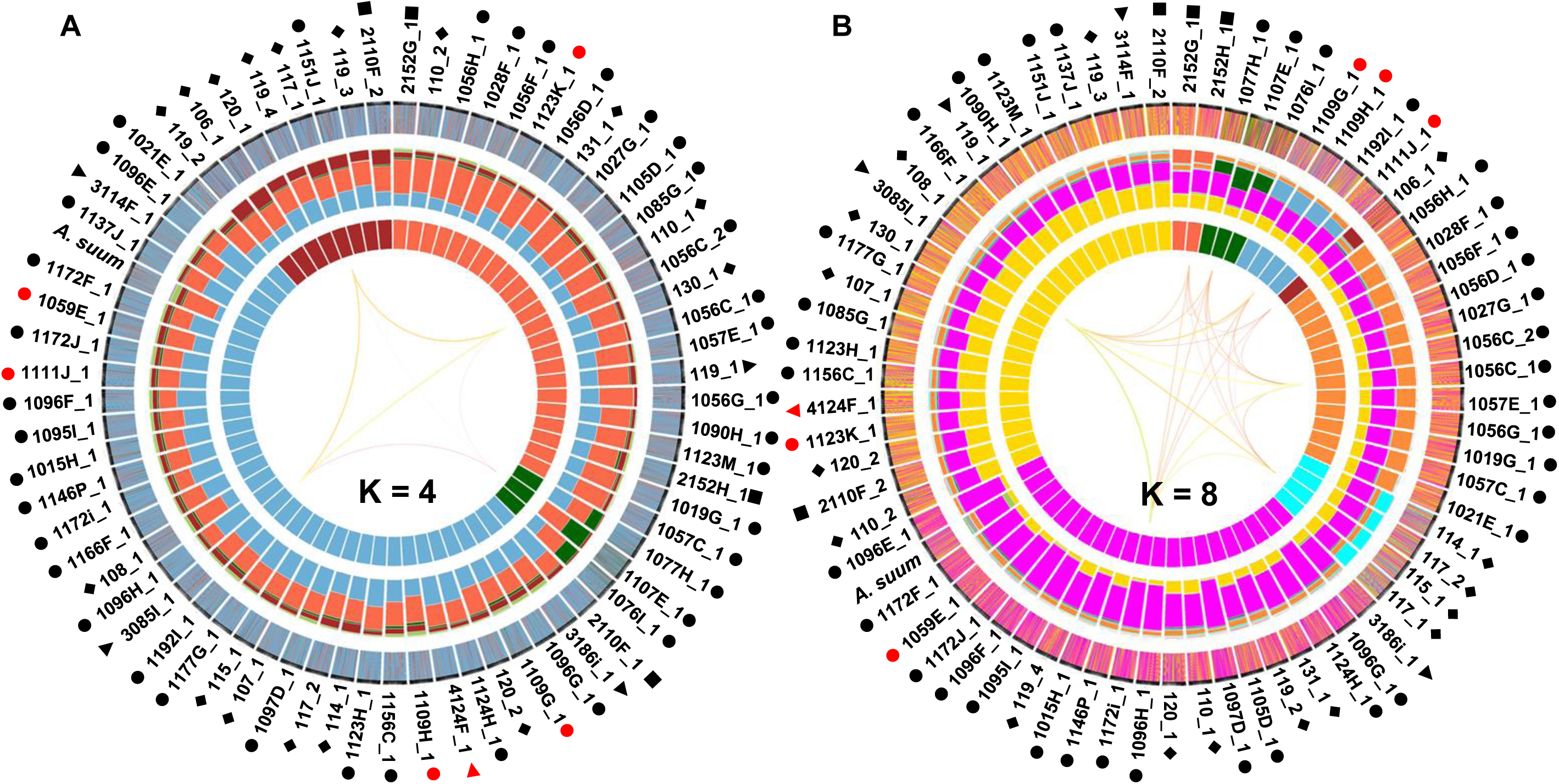
Admixture clustering and current population genetic structure of Ascaris were determined. Data analyzed with POPSICLE with an ancestral population size = 4 (A) and 8 (B) in 10 kb sliding windows as described in Figure 4E.

**Figure S7.**
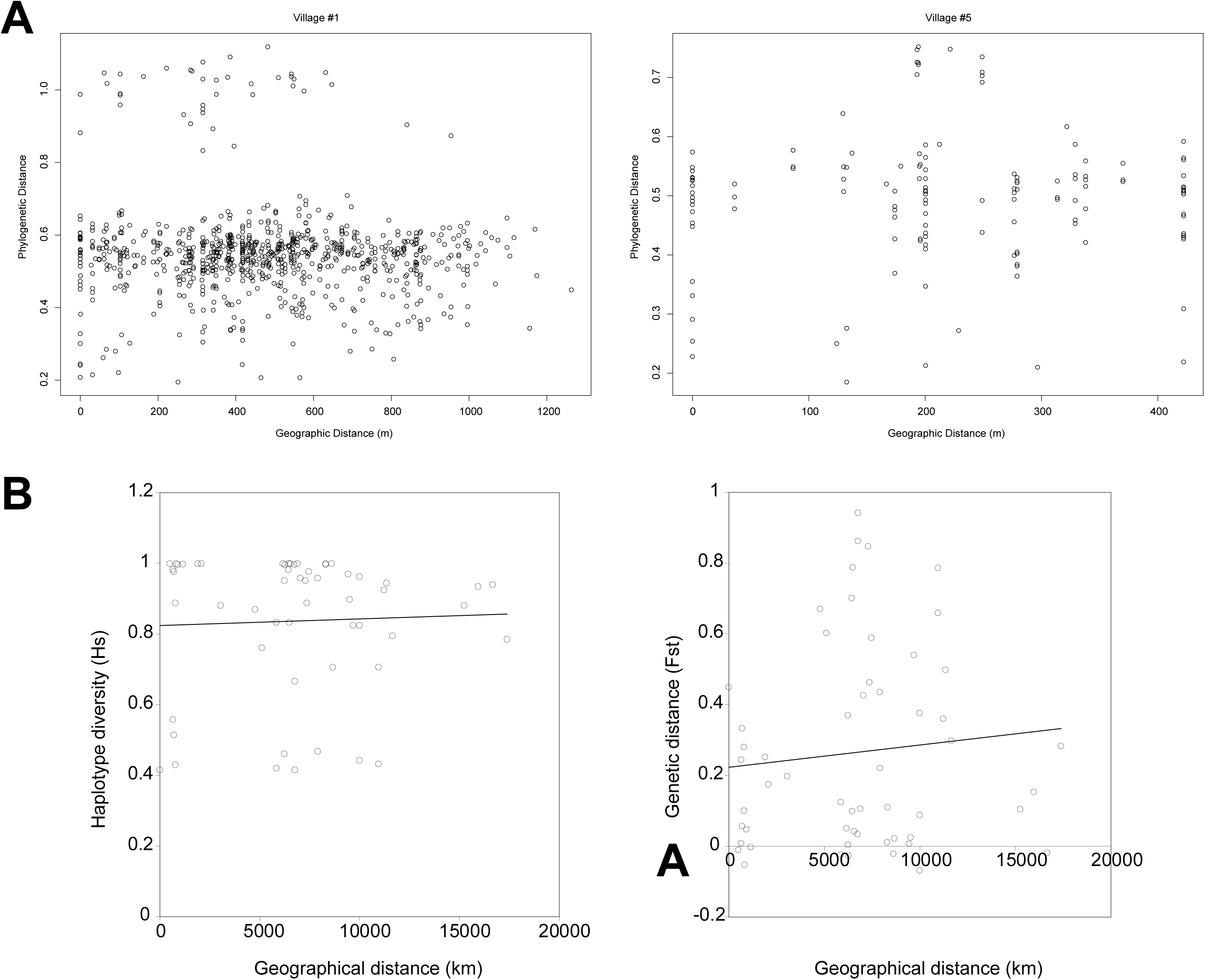
Plot of phylogenetic distances compared to geographic distances. A) For village #1 and village #5. B) Plot of diversity versus geographical distance (Hs on left, Fst on right). Genetic distances based on CO1 genes are plotted against the geographic distances between the places from which these worms were collected.

**Figure S8.**
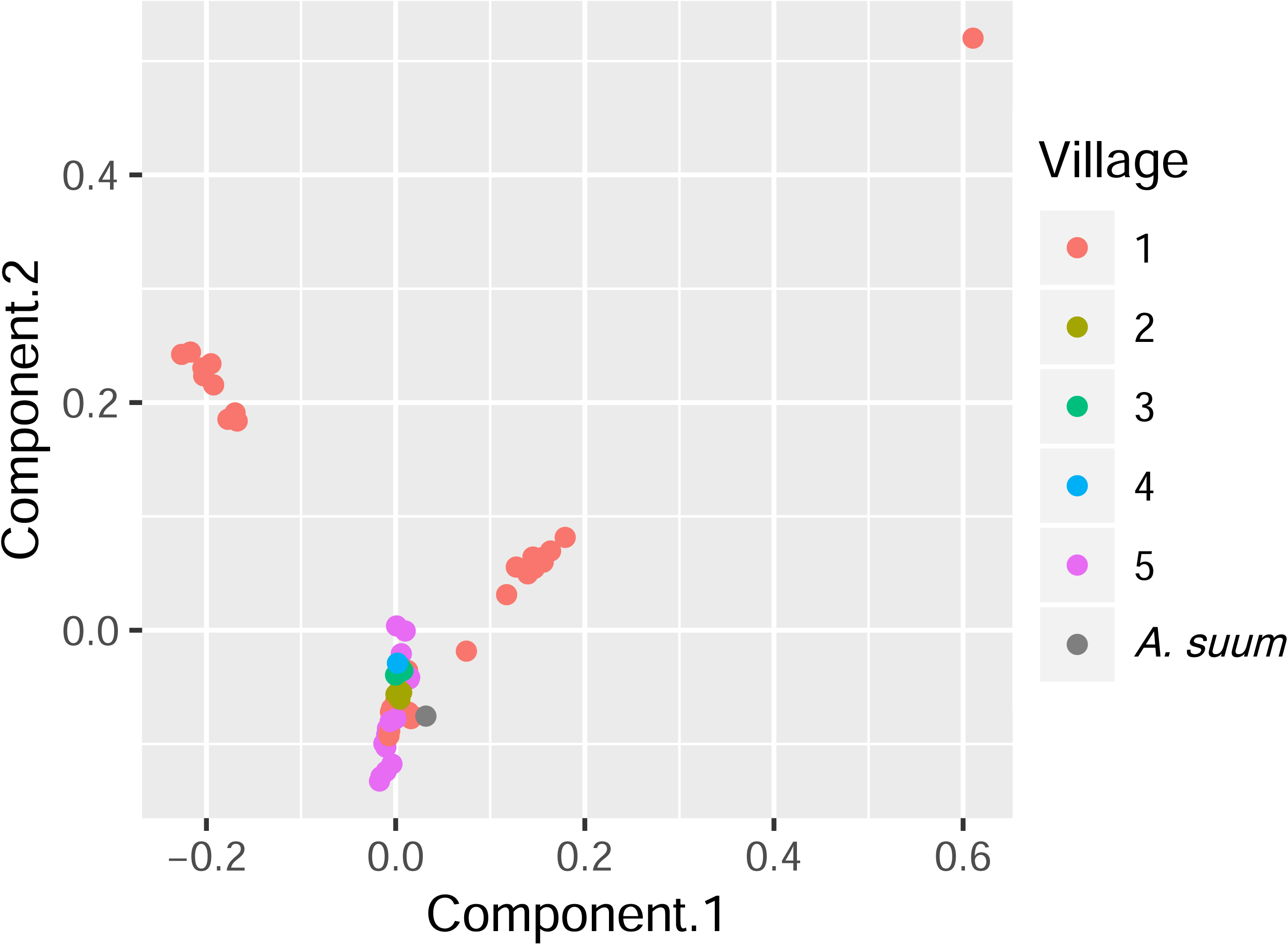
PCA plot of worms sequenced for 5 Kenyan villages. Each point is color-coded by village-of-origin and plotted according to the 1^st^ and 2^nd^ principal components, based on genome sequences. Worms from village #1 are found in each of 3 clusters, and two clusters contain only worms from village #1.

**Figure S9.**
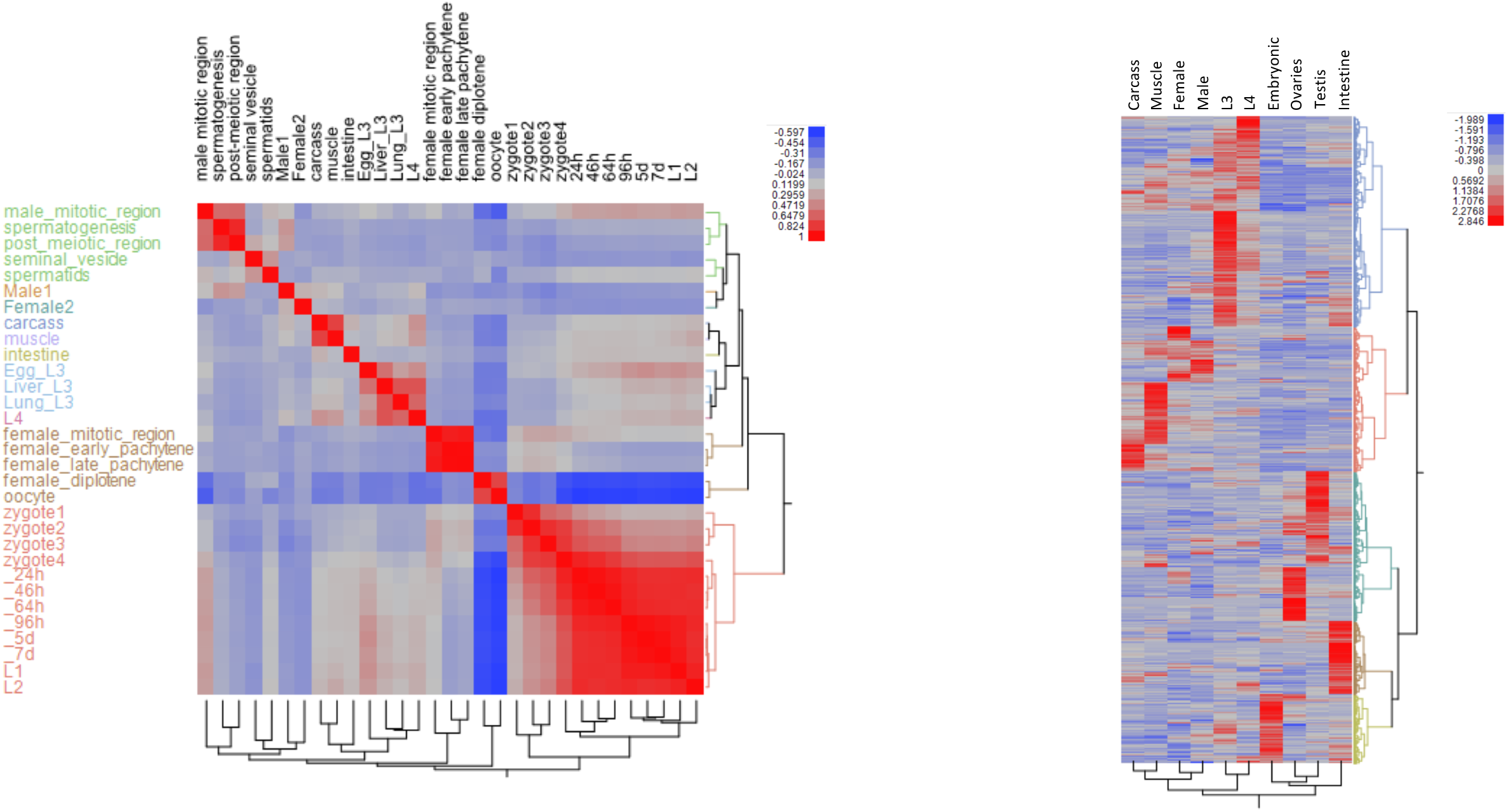
*Ascaris* stage-specific RNA expression heatmaps. (A) Correlation heatmap comparing parasite transcriptomes at different life stages. (B) 1870 genes differentially expressed across the stages.

**Figure S10.**
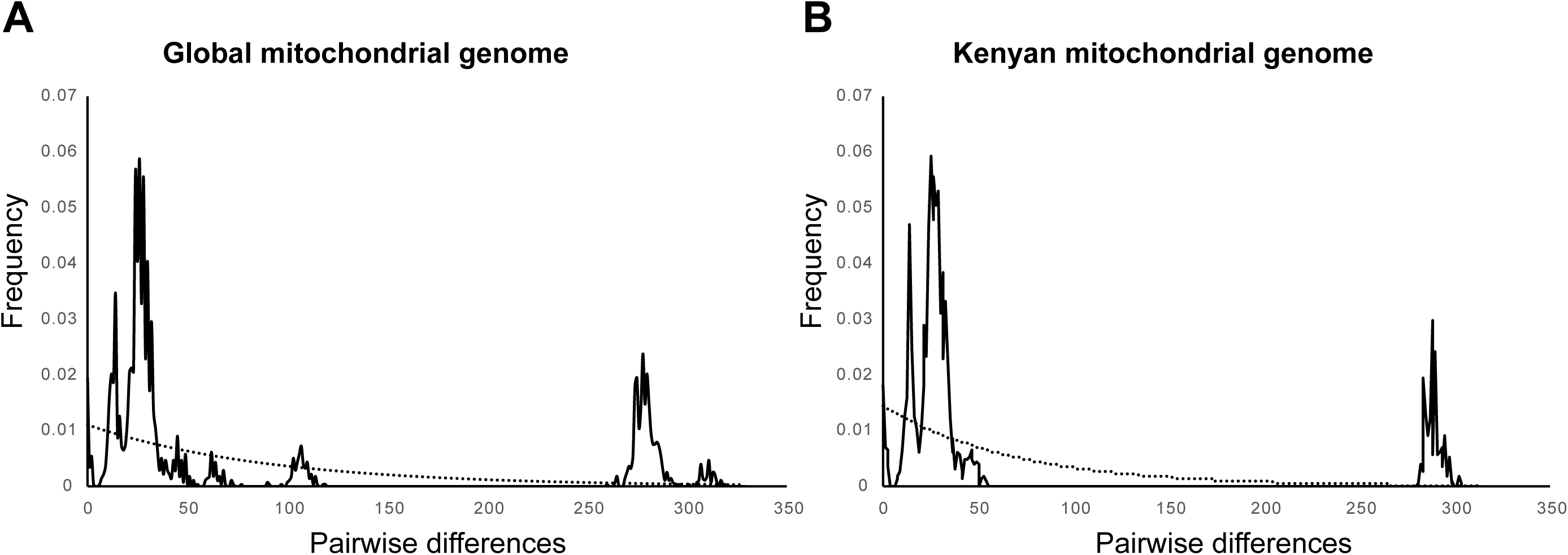
Evidence of *Ascaris* population expansion. The pairwise nucleotide differences between worm samples (solid line) are compared to the binomial function that would most closely represent a theoretical stable population (dotted line). Additional information is available in Table S8.

## Supplementary text

### *Ascaris lumbricoides* reference genome

We first evaluated a *de novo*-only assembly strategy. To facilitate scaffolding, we obtained Pacific Biosciences (PacBio) reads from the female germline (ovary and oviduct) and intestinal tissues. Despite the high coverage of the Illumina paired-end and mate-pair reads and additional longer PacBio reads, the *de novo*-only assembly approach led to a low-quality draft genome with a small N50 value and many scaffolds. We next used a *de novo* assembly strategy with scaffolding to the reference *A. suum* genome (semi-*de novo* strategy). While this assembly led to relatively high N50 values, the assembly contained ∼23 Mb of sequence gaps and an additional 15.4 Mb of sequence that could not be incorporated into the assembly. We believe the reference-based *ALV5.* assembly describe in the main text is the highest quality assembly among the three versions (Table 1) and it should be used for future studies. However, we note that several caveats should be considered when using this genome assembly: (1). potential assembly errors in the reference *A. suum* genome could be propagated into the *ALV5* genome; (2) potential sequence differences between AlV5 and *A. suum*, particularly large-scale genome rearrangements and structure variations, are not likely incorporated into the *ALV5* assembly; and (3) annotation errors and/or missed annotation, either due to errors in the original *A. suum* annotation or due to the potential different gene expression profiles between the two species may be present in the *ALV5* genome. To identify assembly errors and structure variations, we have used additional Illumina sequencing reads to further improve the genome assembly using Pilon genome polishing software. Pilon uses a local re-assemble strategy to correct draft assembly and to call sequence variants of multiple sizes, including very large insertions and deletions ^1^. Overall, the *ALV5* genome assembly is reference-quality and is suitable for downstream analysis, including SNPs and InDels identification. All genome assemblies are made available (see section *Data availability*). Optical mapping, long sequencing reads, and three-dimension chromosomes organization data (Hi-C) will be necessary to further improve the assembly.

### Genome annotation and *Ascaris* proteome

In addition to transferring *A. suum* annotations to the reference-based germline *A. lumbricoides* genome, we also transferred the *A. suum* gene models to the *de novo* and semi-*de novo A. lumbricoides* genome versions. While ∼94.6% of the genes can be transferred to both genomes, over 20% of the transferred genes are only partial matches and are fragmented supporting the view that the *de novo* and semi *de novo A. lumbricoides* assemblies are highly fragmented. Although the updated *Ascaris suum* proteome was reported in our previous study ^2^, that study’s focus was on comparative analysis of programmed DNA elimination. Thus, no in-depth analysis of the revised *Ascaris* proteome (based on the annotation of the reference-based genome) was carried out in that study.

### Nucleotide diversity and population structure inferred from mitochondrial genomes

Sliding window analyses showed considerable diversity across *Ascaris* mitochondrial genomes. The most diverse region was the non-coding control region, both between worms from different villages in this study and between Kenyan and other worms. The ND4 gene had a nucleotide diversity of π = 0.008827; that of the CO1 barcoding gene was 0.006243 (Figure S2). Despite differences in their molecular diversity, phylogenetic analyses based on the ND4 and CO1 genes revealed the same overall topology: distinct *A. lumbricoides* and *A. suum* type clades (Figure S3). Though the phylogeny passed on the whole mitochondrial genomes also suggests the presence of an *A. lumbricoides*-type clade and an *A. suum*-type clade, the mitochondrial genome generated from the Sanger institute *A. suum* genome reads (Acc: PRJEB2435) clustered into the *A. lumbricoides* type clade (Figure 3B).

### Evolution of *Ascaris* mitochondrial genomes

To account for potential population expansion events, both datasets including all published mitochondrial genomes from across the globe and those from Kenya alone were assessed (Figure S10 and Table S8). The Tajima’s D value was negative and significant (Tajima’s D −1.5691; P-value 0.028, P<0,05) indicating an excess of low frequency polymorphisms within the global population data set. The Fu’s *Fs* for the global data was shown to be positive but not significant (Fu’s *Fs* 8.5673; P-value 0.98, P>0.05), potentially indicating a deficiency in diversity as would be expected in populations that has recently undergone a bottle neck event. Furthermore, the same patterns were also seen when only the Kenyan sequences were compared to each other where the Fu’s *F*_*S*_ was positive and nonsignificant (Fu’s *F*_*S*_ 4.979; P-value 0.917, P>0.05), and the Tajima’s *D* ^43^ was also negative but nonsignificant (Tajima’s *D* −1.28930; P-value 0.079, P>0.05.

Overall, these data suggest that *Ascaris* either underwent a population expansion or a significant selective pressure. To test for selection on the *Ascaris* population, we calculated the ratio of non-synonymous to synonymous substitutions (dN/dS ratio) for each of the genes in the mitochondrial genome. All genes appeared to be under purifying selection with a dN/dS ratio <1 (with ND6 having the lowest ratio value of 0.05 and ND5 with the highest ratio value of 0.527). Since we observed a dN/dS ratio below 1 for all genes, it is unlikely that the genetic variability of the *Ascaris* samples examined here resulted from selective sweeps. This means that these genes are likely under purifying selection to maintain the existing functionality of these mitochondrial-encoded proteins, as opposed to drastic changes resulting from selective sweeps. Thus, the Tajima’s *D* and Fu’s *F* measurements are most likely due to a population expansion. Since we do not know how quickly *Ascaris* mutates, we cannot estimate how recently such a population expansion might have occurred. Results were similar when based on sequences of worms collected only in Kenya, suggesting that a recent (worldwide) *Ascaris* population expansion may have occurred.

### Population structure inferred from nuclear genomes

Most clades in the nuclear genome phylogeny are dominated by worms from either village #1 or village #5, which were the most heavily parasitized and also the most heavily represented in this sampling. P-values for clustering of worms from the same host-groups (Table 2) are shown with Bonferroni corrections for multiple comparisons. The two significant groupings (for the nuclear phylogeny) were individual and village. These remain significant if both factors are included in the same regression, or if the “individual” variable is treated as being nested within the “village” variable.

Geographic distance was not a significant predictor of phylogenetic distance across the five villages (r2=0.07, p=0.15) within village #5 (r^2^=0.05, p=0.27), or within village #1 (r^2^=-0.07, p=0.81). This analysis was only done for these two villages, as they had the largest number of worms sequenced in this study (Figure S6).

## Supplementary Methods

### DNA extraction from adult worms

DNA extraction methods were modified several times before a protocol was identified that produced DNA of sufficient quality. Extraction was likely difficult because the worms have a high polyphenol content ^3^. In the end, approximately 200-400 mg of worm material was put in a 1.5 mL tube (Eppendorf, Hamburg, Germany). To the sample, 750 μl of Qiagen Digestion buffer G2 (Qiagen, Hilden, Germany) and 20 μl of Proteinase K (Qiagen, Hilden, Germany) was added. Tubes were manually shaken for ten seconds, and then incubated in a Thermomixer C heat block (Eppendorf, Hamburg, Germany) at 50 degrees and 500 rpm for four hours. Halfway through the incubation, tubes were manually shaken again. Tubes were spun at 10000 rpm for one minute, and 700 μl of the supernatant was removed and to which 700 μl one-phase Phenol/Chloroform/Isoamyl alcohol (Amresco LLC, Solon, Ohio, USA) was added to these tubes. Tubes were inverted 30 times, then spun 4°C at 14000 rpm for 20 minutes. The supernatant was removed, and 3 μl RNase A (ThermoFisher Scientific, Waltham, MA, USA) was added. The sample was mixed and incubated at 37°Cfor 30 minutes at 200 rpm. Once again, 700 μl one-phase Phenol/Chloroform/Isoamyl alcohol was added to these samples, the samples were inverted 30 times, and spun for 15 minutes. The supernatant was then removed and added to the Zymo Genomic DNA Clean & Concentrator kit.

### Analysis of mitochondrial genomes

Genetic diversity was compared to the CO1 gene, as the CO1 gene has been used extensively in molecular epidemiological studies of *Ascaris* and other nematodes. The sliding window was performed on all mitochondrial genomes and the subset containing only those from Kenya. For comparison, sliding window analyses were also performed on the complete CO1 gene in order to identify if the barcoding region itself was actually the most diverse region of the gene to be used.

Non-coding and tRNA sequences were removed from the analyses, to remove the potential effect of high mutation rates which could have caused mutational saturation in any of the analyses and masked the true impact of the divergence in the protein coding genes.

## Supplementary Tables

**Table S1. Characteristics of genome assemblies**

**Table S2. Proteome annotation**

**Table S3. Description of worm from which each sample was sequenced.** The sex of the worm (based on morphological identification) and the part of the worm (germline vs somatic) is listed. Some hosts donated multiple worms.

**Table S4. CO1 haplotype list**

**Table S5. X4 ratio analyses of Clade A and B using complete mitochondrial genomes used to construct the phylogeny in Fgure 2b.**

**Table S6. Number of heterozygous and homozygous SNPs in each of the 68 worms from Kenya sequenced.**

**Table S7. Reference mitochondrion genomes**

**Table S8. Demographic analyses using Tajima’s D and Fu’s F statistic across complete mitochondrial genomes as a detection for the signature of population expansion events.**

**Table S9. Supplement to Table 2 using alternative measures of phylogenetic distance**

## Supplementary figure legends in main manuscript file

## Notes

### Competing Interest Statement

The authors have declared no competing interest.

