## Supplemental Table 5 for "Extensive hybridization between pig and human *Ascaris* identifies a highly interbred species complex infecting humans"

**Table S5. X4 Ratio analyses of Clade A and B using complete mitochondrial genomes using to construct the phylogeny in figure 2b**

| **parameter** | **Cluster 1** | **Cluster 2** |
| --- | --- | --- |
| n | 66 | 18 |
| k | 0.0235 | 0.0235 |
| d | 0.01 | 0.01 |
| π =(d*(n/n-1) | 0.01015 | 0.010588 |
| ϴ = π/(1-(4π/3) | 0.010289 | 0.01074 |
| k/ϴ | 2.283938 | 2.188111 |

Where n= numbers of sequences; K=total pairwise differences between clades; d = pairwise differences within clades
