## Supplemental Table 8 for "Extensive hybridization between pig and human *Ascaris* identifies a highly interbred species complex infecting humans"

| Population | No. Sep | No. haps | Tajima’s D |  | Fu’s F |  |
| --- | --- | --- | --- | --- | --- | --- |
| Global | 85 | 48 | -1.5691 | (P 0.028) | 8.5673 | (P 0.975) |
| Kenya | 68 | 35 | -1.28930 | (P 0.079) | 4.979 | (P 0.917) |

Table S7. Demographic analyses using Tajima’s D and Fu’s F statistic across complete mitochondrial genomes as a detection for the signature of population expansion events
